# Novel compounds derived from AR-12 that demonstrate host-directed clearance of intracellular *Salmonella enterica* Serovar Typhimurium

**DOI:** 10.64898/2025.12.30.696991

**Authors:** Elizabeth G. Graham-Gurysh, M. Shamim Hasan Zahid, Devika M. Varma, Antonio Landavazo, Ojas A Namjoshi, Joseph W Wilson, Monica M. Johnson, Ryan N. Woodring, Aaron T. Hendricksen, Joseph Vath, Erica N. Pino, Eric M. Bachelder, Bruce E. Blough, Kristy M. Ainslie

## Abstract

*Salmonella* infections, including typhoid and paratyphoid fevers, impose a substantial burden in low-income countries, contributing to significant morbidity and mortality associated with enteric and systemic infections. Developing effective treatments for *Salmonella* infections requires further consideration, especially due to widespread antibiotic resistance. Host-targeted therapies, such as AR-12, offer a promising solution to combat drug resistance. AR-12 has demonstrated broad-spectrum antimicrobial activity against various bacterial pathogens, including *Salmonella enterica* serovar Typhimurium, *S*. Typhi, *Francisella tularensis*, and *F. novicida*, as well as protozoan parasites and fungal pathogens. To expand its HDT potential against *S. Typhimurium*, we conducted a medicinal chemistry campaign using AR-12 as a scaffold with systematic optimization of various points of diversity and the pyrazole core to develop analogs informed by structure–activity relationship. This work led to the development of 81 AR-12 analogs. Primary screening identified 38 analogs that are both more potent and less cytotoxic than parent compound AR-12, while only three were less potent and more cytotoxic. Further, only seven compounds affected planktonic *Salmonella* growth below 20 µM, suggesting host-directed activity in most of the compounds. Twelve analogs were chosen for secondary screening in MDR *S*. Typhimurium. Compounds 372, 373, and 378 demonstrated remarkable selectivity, with values exceeding 1500 for both susceptible and MDR *S*. Typhimurium, compared to AR-12’s selectivity of around 20. This approximately 100-fold improvement, coupled with improved potency against intracellular *Salmonella*, suggests these analogs have significantly greater host-directed activity than direct antibacterial effects. Proteomic analysis for the two most potent compounds, 341 and 370 revealed enrichment of vesicle-mediated transport proteins, specifically with respect to retrograde transport at the trans-Golgi-network and intra-Golgi traffic. These results suggest that the analogs reduce intracellular *S*. Typhimurium replication by disrupting its exploitation of the host cell’s vesicle-mediated transport system.

## INTRODUCTION

Salmonellosis is a major bacterial enteric illness caused by contaminated food and water that can result in death and irreparable gastrointestinal damage. According to the WHO, the causative agent of salmonellosis, *Salmonella*, is responsible for one in four cases of diarrheal diseases worldwide.(1) *Salmonella* infection presents as typhoid and paratyphoid fever, which is caused by gram-negative bacteria *Salmonella enterica* serovars Typhi and Paratyphi (A, B, C), respectively.(2) In the US, non-typhoidal *Salmonella* strains alone cause 1.35 million infections per year amounting to $400 million in direct medical costs.(3) While *S*. Typhi uses human host as its exclusive reservoir, *S*. Typhimurium can be isolated from human, animal, and retail meat origins worldwide and are largely responsible for non-typhoidal cases of *Salmonella* infections.(4, 5)

Salmonellosis affects people globally, but typhoid and paratyphoid fevers predominantly impact young children in low-income countries. Therefore, when developing treatments for *Salmonella* infections, it is crucial to carefully consider factors like toxicity. The widespread antibiotic resistance of *S. enterica* to first and second-line treatments has led to a reliance on fluoroquinolones, which are associated with higher toxicity.(20) For patients who do not respond to fluoroquinolones like ciprofloxacin, alternative combinations such as azithromycin or aztreonam are used. Research indicates that the fatality rate for hospitalized typhoid patients can increase from 1-2% to 10-20% if inappropriate antimicrobials are used. This issue is particularly challenging in developing nations with limited diagnostic capabilities.(21)

The overuse or misuse of antibiotics over several decades has been the primary cause for the development of antibiotic resistance in *S. enterica* serovars. In recent decades, clinical isolates of *S*. Typhi have illustrated large-scale resistance against first-line antibiotics (chloramphenicol, ampicillin, and trimethoprim-sulfamethoxazole) and are now termed multi-drug resistant (MDR) strains. Treatment options are drastically tapering amidst reports of extensively drug resistant isolates of *S*. Typhi in Asia which are resistant to first, second and third-line antibiotics, including cephalosporins.(6) Drug resistance is increasingly reported even in non-typhoidal cases of the diseases with ∼10% being resistant to ciprofloxacin, a commonly available second-line fluoroquinolone used to treat enteric fever.(3) Adding to this, there is a general dearth of novel antibiotics against gram negative pathogens due to the slowing of the antibiotic pipeline over the last decade and existing last line therapies suffer from toxicity concerns.(7) Thus, new treatment options that are effective against resistant strains of *S. enterica* are needed.

Targeting the host over the pathogen directly is one method to combat the emergence of drug resistance.(8, 11, 22, 23) Host directed therapies (HDTs) that target host-encoded functions necessary for bacterial infection, replication, virulence, and pathogenesis have the potential to be more effective than direct antimicrobials in treating intracellular infections.(8) Most importantly, HDTs can counter the larger problem of antimicrobial resistance by targeting host cells, such as macrophages and intestinal epithelial cells, relieving the direct selective pressure on bacteria that typically drives drug resistance.(9, 10) Several novel and repurposed drugs are being evaluated for host-directed therapies against bacterial infections.(9, 11) AR-12 (originally named OSU-03012) is one such IND-FDA approved small molecule, initially developed as a chemotherapeutic agent. AR-12 has demonstrated impressive broad-spectrum host-targeted antimicrobial activity. It has been effective against a range of bacterial pathogens—including *Salmonella Typhimurium, (13, 24-27) S*. Typhi,(28) *Francisella tularensis* (Schu S4 and LVS), and *Francisella novicida* (12, 29)—as well as the protozoan parasite *Leishmania donovani* and *L. mexicana* (14, 30, 31) and the fungal pathogen *Cryptococcus neoformans*.(32, 33) Remarkably, AR-12 also exhibits direct antiviral activity against high-consequence pathogens such as Lassa, Ebola, Marburg, and Nipah viruses.(34) Moreover, when used in combination with the FDA-approved drug sildenafil, AR-12 effectively downregulates multiple host receptors involved in viral infection.(35) We have also shown that AR-12 can enhance the activity of ampicillin and streptomycin in resistant intracellular *S*. typhimurium.(27)

Despite its broad host-directed antimicrobial activity, AR-12 has a narrow therapeutic window for *Salmonella* (IC□□ = 0.4 µM; LC□□ = 8 µM) which limits its translational potential. To expand its HDT potential, we conducted a medicinal chemistry campaign using AR-12 as a scaffold to develop analogs informed by structure–activity relationship to identify improved activity against *S. Typhimurium*. We screened 81 AR-12 analogs (Supplemental Table 1) for activity against *S*. Typhimurium (Figure 1). Primary screening of 81 AR-12 analogs assessed changes in intracellular *S*. Typhimurium burden (IC_50_), host cell viability (LC_50_), and direct effects on planktonic *S*. Typhimurium (MIC_50_), calculating selectivity (LC_50_/IC_50_) to determine host-directed potency versus cytotoxicity. From this, we identified 12 hit compounds with high selectivity or high potency that are less cytotoxic to the host cell than the parent compound AR-12. These compounds activity were then investigated against MDR *S*. Typhimurium and the two most potent compounds were evaluated by proteomic analysis.

**Figure 1:**
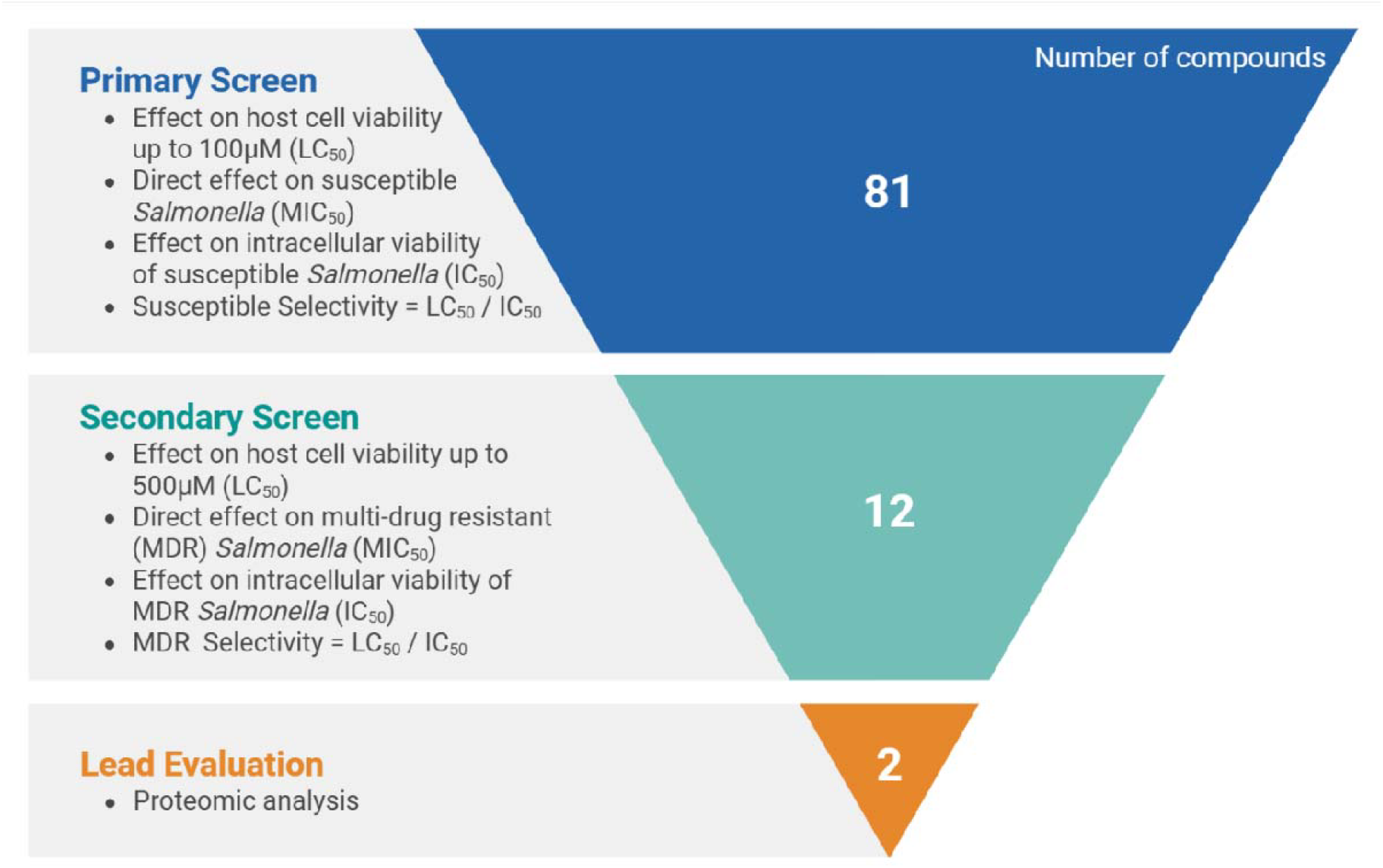
Process flow of screening methodology and hit selection. Primary screening: compounds evaluated against susceptible (Susc.) *S*. Typhimurium. Compounds with selectivity > 250 or extremely potent IC_50_ <0.1µM while being less cytotoxic than AR-12 were identified as hits for secondary screening against multi-drug resistant (MDR) *Salmonella*. The two most potent compounds against MDR *Salmonella* underwent proteomic analysis.

## RESULTS AND DISCUSSION

Using AR-12 as a scaffold, we conducted a medicinal chemistry campaign to optimize the scaffold for pre-clinical development, focusing on enhancing potency against intracellular infection while minimizing direct effects. Our strategy involved systematically optimizing various points of diversity and the pyrazole core which helped us to develop structure-activity relationship (SAR) models to reduce macrophage bacterial burden while also decreasing cytotoxicity to host cell (**Figure 2**). Our medicinal chemistry campaign used an iterative approach, optimizing multiple points of diversity on the molecule, simultaneously.

**Figure 2:**
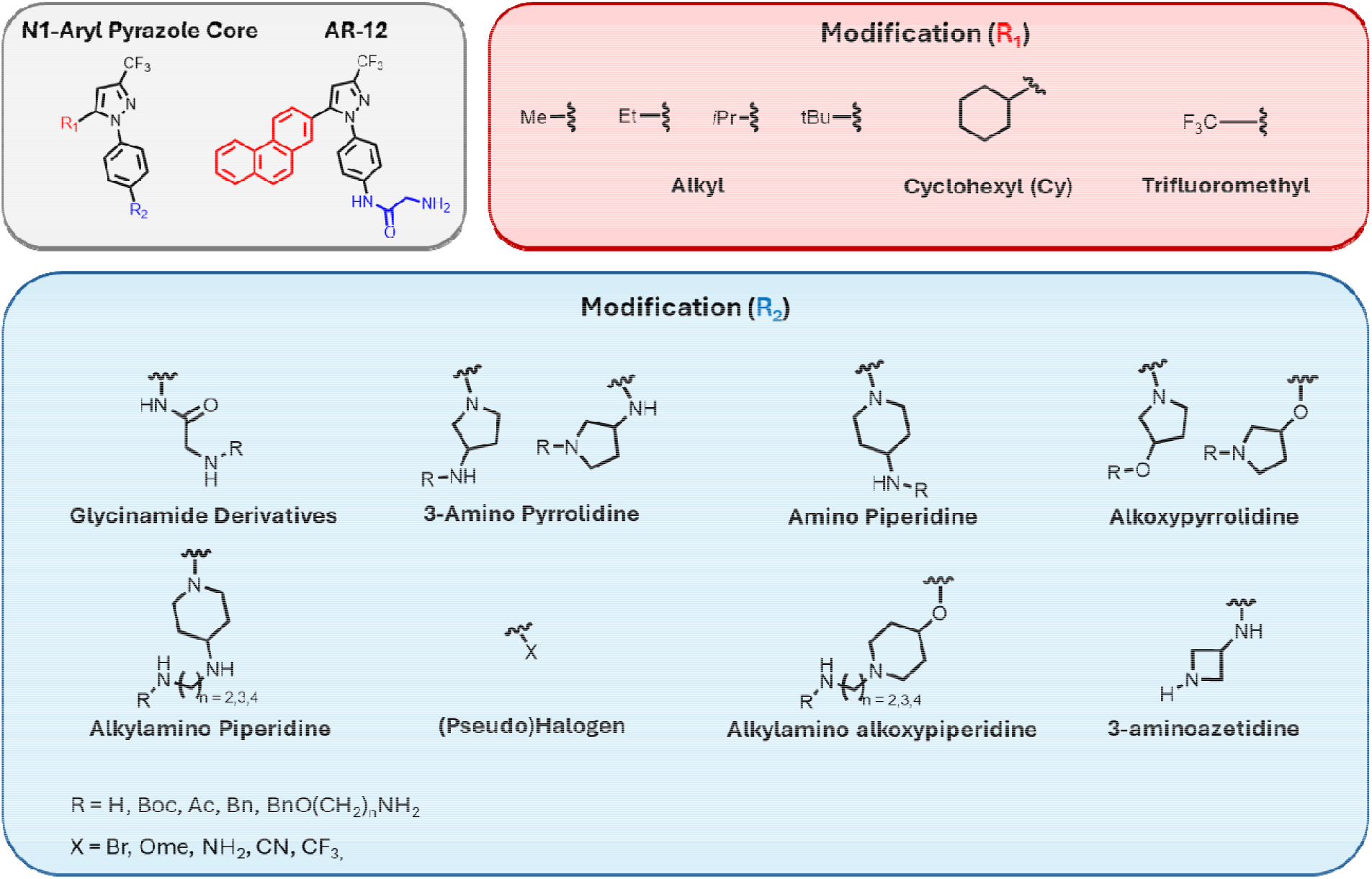
Novel AR-12 analogs with host-directed clearance of intracellular *Salmonella enterica* Serovar Typhimurium. AR-12 contains an N1-aryl-3-trifluoromethyl pyrazole core with variable R□ (red) and R□ (blue) positions. A library of 81 analogs was synthesized with systematic R□/R□ substitutions and screened for host-directed antibacterial activity using a gentamicin protection assay, effect on host cell viability, and direct effect on planktonic *S*. Typhimurium. R□ modifications were primarily small alkyl groups (methyl, ethyl, isopropyl, tert-butyl, cyclohexyl), while R□ incorporated diverse polar, basic, and electronegative motifs— including glycinamide derivatives, 3-amino pyrrolidine, amino piperidine, alkoxypyrrolidine, alkylamino piperidine, halogens and pseudohalogens (CN, CF□, OMe, NH□), and 3-aminoazetidine.

### Primary Screen

Primary screening of all 81 AR-12 analogs (**Supplemental Table 1**) assessed the change in intracellular *Salmonella* burden (IC_50_), host cell viability (LC_50_), and direct effect of planktonic *Salmonella* (MIC_50_) after treatment with compounds (**Table 1**). From this data, selectivity was calculated (LC_50_ / IC_50_) to determine the relative potency of the host-directed effects compared to host cell toxicity. Interestingly, only seven analogs had any effects on planktonic *Salmonella* growth below 20 µM. Additionally, for those compounds with some direct effects, the IC_50_ was between 10 to 140-fold lower than the MIC_50_ (**Supplemental Table 2**). This strongly suggests that the analogs act in a host-directed manner and would be unlikely to exert direct pressure that might drive drug resistance. Of the 81 compounds, 38 were both less cytotoxic and more potent than parent compound AR-12, while only 3 were less potent and more cytotoxic (**Supplemental Figure 1**). From these results we identified twelve hit compounds (**Figure 3**) with high selectivity (>250) or high potency (IC_50_ < 0.1µM) that were less cytotoxic than AR-12. These hit compounds were selected for secondary screening in multi-drug resistant (MDR) *Salmonella*.

**Table 1:** Results of primary screen. Concentration at which intracellular susceptible *S*. Typhimurium burden is 50% in RAW 264.7 macrophages determined by gentamicin protection assay (Susc. IC_50_) and concentration where RAW 264.7 macrophage cell viability is reduced by 50% (LC_50_) as determined by MTT assay. The concentration of compounds that reduce planktonic susceptible *S*. Typhimurium viability to 50% as measured by optical density (MIC_50_). Parental compound AR-12 is included for comparison. Compounds highlighted in light gray were selected for a secondary screen. ND = not determined.

| ID | $\text{IC}_{50}$<br>($\mu\text{M}$) | $\text{LC}_{50}$<br>($\mu\text{M}$) | Select-<br>ivity | $\text{MIC}_{50}$<br>($\mu\text{M}$) |
| --- | --- | --- | --- | --- |
| AR-12 | 0.42 | 8.3 | 19.8 | $>20$ |
| 202 | $>5$ | $>100$ | ND | $>20$ |
| 203 | 0.11 | 20.6 | 192 | $>20$ |
| 229 | $>5$ | $>100$ | ND | $>20$ |
| 230 | $>5$ | $>100$ | ND | $>20$ |
| 232 | 0.28 | 30.6 | 109 | $>20$ |
| 247 | 0.20 | 7.0 | 35 | $>20$ |
| 285 | 0.34 | 56.2 | 165 | $>20$ |
| 286 | 0.61 | 41.7 | 68 | $>20$ |
| 312 | 0.13 | 11.1 | 83.8 | $>20$ |
| 313 | 0.25 | 60.9 | 244 | $>20$ |
| 314 | 0.13 | 27.2 | 214 | $>20$ |
| 315 | 0.16 | 28.6 | 185 | $>20$ |
| 316 | $>5$ | $>100$ | ND | $>20$ |
| 317 | $>5$ | $>100$ | ND | $>20$ |
| 318 | 0.20 | 30.1 | 152 | $>20$ |
| 319 | 0.19 | 6.6 | 35 | $>20$ |
| 321 | 0.97 | 25.4 | 26 | $>20$ |
| 322 | 0.14 | 43.7 | 324 | $>20$ |
| 323 | 0.10 | 32.0 | 333 | $>20$ |
| 324 | 0.08 | 18.8 | 244 | $>20$ |
| 327 | 2.97 | $>100$ | $>34$ | $>20$ |
| 330 | 0.76 | 56.0 | 74 | $>20$ |
| 334 | 0.27 | 30.1 | 112 | $>20$ |
| 336 | 0.30 | 28.5 | 95 | $>20$ |
| 337 | 0.37 | 29.4 | 80 | $>20$ |
| 338 | $>5$ | 47.3 | $<9$ | $>20$ |
| 339 | 0.45 | 29.1 | 65 | $>20$ |
| 340 | $>5$ | 37.2 | $<7$ | $>20$ |
| 341 | 0.08 | 31.8 | 398 | $>20$ |
| 352 | 0.65 | 29.1 | 45 | $>20$ |
| 353 | 0.33 | 26.4 | 80 | $>20$ |
| 354 | 0.98 | 40.3 | 41 | $>20$ |
| 355 | 0.22 | 20.3 | 92 | $>20$ |
| 356 | 0.22 | 24.7 | 113 | $>20$ |
| 357 | 0.35 | 28.5 | 82 | $>20$ |
| 358 | 0.55 | 41.7 | 76 | $>20$ |
| 362 | $<0.10$ | 6.2 | $>62$ | $>20$ |
| 363 | 0.98 | 57.2 | 58 | $>20$ |
| 364 | 0.13 | 35.0 | 269 | $>20$ |
| 365 | 0.15 | 42.7 | 285 | $>20$ |
| 366 | 1.70 | 34.0 | 20 | $>20$ |
| 367 | $>5$ | $>100$ | ND | $>20$ |
| 368 | $<0.10$ | 6.5 | $>65$ | $>20$ |

| ID | IC<sub>50</sub><br>(μM) | LC<sub>50</sub><br>(μM) | Select-<br>ivity | MIC<sub>50</sub><br>(μM) |
| --- | --- | --- | --- | --- |
| 370 | 0.08 | 17.2 | 229 | >20 |
| 371 | 0.14 | 90.2 | 644 | >20 |
| 372 | 0.13 | >100 | >769 | >20 |
| 373 | 0.13 | >100 | >769 | >20 |
| 374 | 0.14 | 3.2 | 23 | >20 |
| 375 | 0.23 | 29.5 | 128.1 | >20 |
| 376 | 0.47 | >100 | >212 | >20 |
| 377 | 0.39 | 31.4 | 81 | >20 |
| 378 | 0.31 | >100 | >322 | >20 |
| 381 | 0.11 | 28.6 | 260 | >20 |
| 389 | 0.60 | >100 | >167 | >20 |
| 392 | 0.56 | 3.5 | 6 | >20 |
| 394 | 0.23 | 6.8 | 29 | >20 |
| 395 | 0.28 | 11.3 | 40 | >20 |
| 396 | 0.11 | 17.1 | 157 | >20 |
| 397 | 0.62 | 23.2 | 37 | >20 |
| 412 | >5 | >100 | ND | >20 |
| 413 | 0.71 | >100 | >141 | >20 |
| 414 | >5 | >100 | ND | >20 |
| 415 | >5 | >100 | ND | >20 |
| 416 | 0.21 | 8.1 | 38.5 | 17 |
| 417 | 0.40 | 8.7 | 22 | >20 |
| 418 | 0.12 | 26.4 | 225 | 16 |
| 419 | 0.30 | 9.2 | 31 | >20 |
| 420 | 0.36 | 6.2 | 17 | 17 |
| 421 | 0.16 | 10.5 | 66 | >20 |
| 422 | 0.19 | 2.2 | 12 | >20 |
| 423 | 0.26 | 2.3 | 9 | >20 |
| 424 | 0.17 | 1.8 | 10 | 20 |
| 425 | 0.23 | 1.9 | 8 | 11 |
| 426 | 0.19 | 17.8 | 94 | >20 |
| 427 | 0.81 | 31.0 | 38 | >20 |
| 428 | 0.40 | 18.5 | 46 | >20 |
| 429 | 1.33 | 2.2 | 2 | 14 |
| 430 | 0.23 | 26.4 | 115 | >20 |
| 431 | 0.18 | 7.0 | 39 | >20 |
| 432 | 0.51 | 4.6 | 9 | >20 |
| 433 | 0.39 | 8.2 | 21 | 17 |

**Figure 3:**
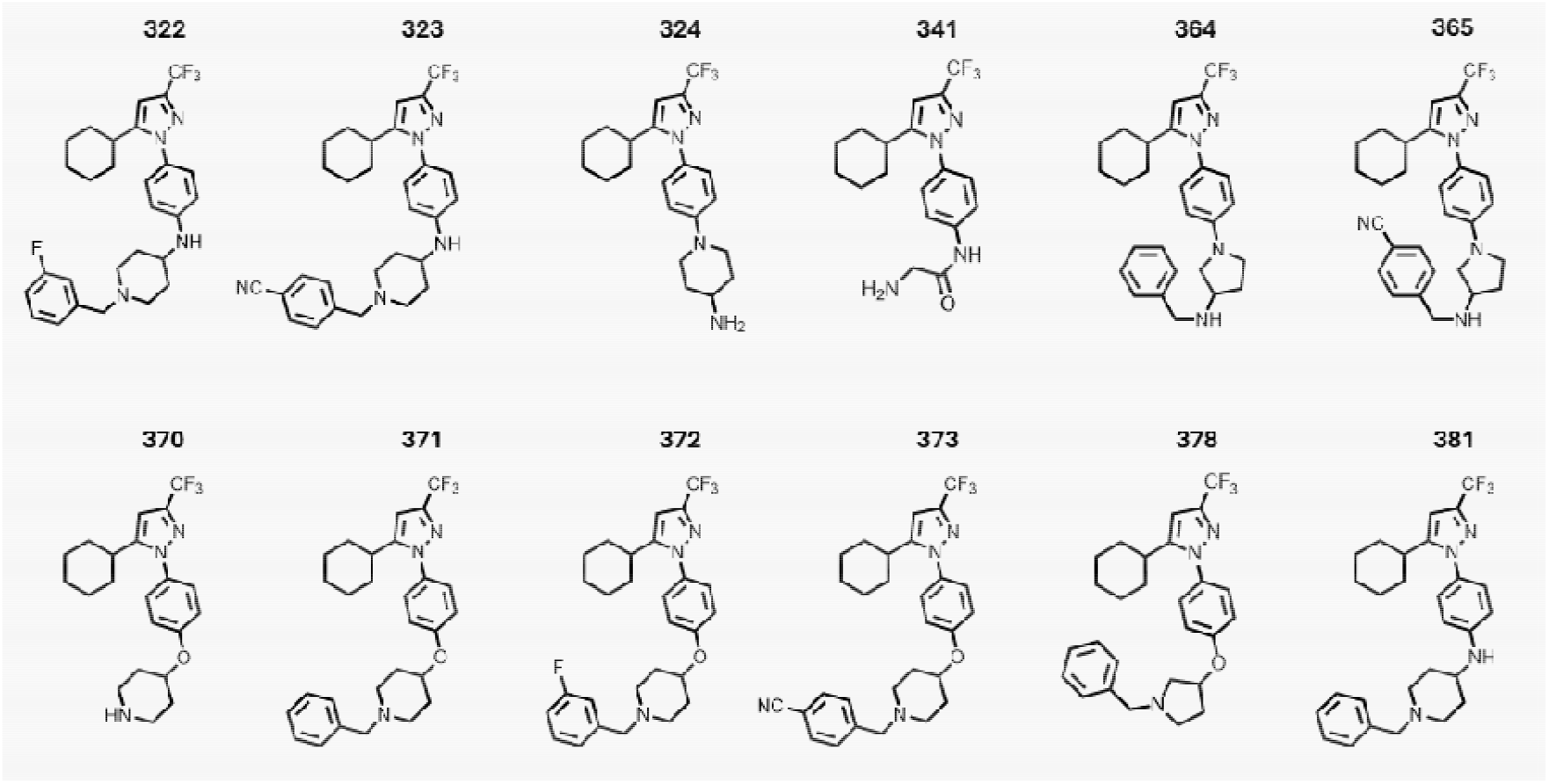
Lead AR-12 analogs. Chemical structure of 12 compounds with high susceptible selectivity (>250) or extremely potent (IC_50_ < 0.1µM) and less cytotoxic than AR-12 were selected for secondary screening.

### Secondary Screen

A secondary screen was performed by assessing intracellular growth of MDR *S*. Typhimurium in RAW 264.7 macrophages by gentamicin protection assay (**Supplemental Figure 2**). The effect of compounds on host cell viability was extended up to 500 µM. Additionally, the direct effect of analogs on growth of MDR *S*. Typhimurium was also investigated (**Table 2**). As with susceptible *S*. Typhimurium, no hit compounds had any direct effect on planktonic bacterial viability up to 20µM. Notably, only 3 compounds (323, 324, 365) had strongly different IC_50_ between susceptible and MDR *S*. Typhimurium (defined as 2-fold difference MDR IC_50_ / Susc IC_50_). The largely agnostic efficacy between susceptible and MDR bacteria further supports the notion that these compounds are acting in a host-directed manner.

**Table 2:**
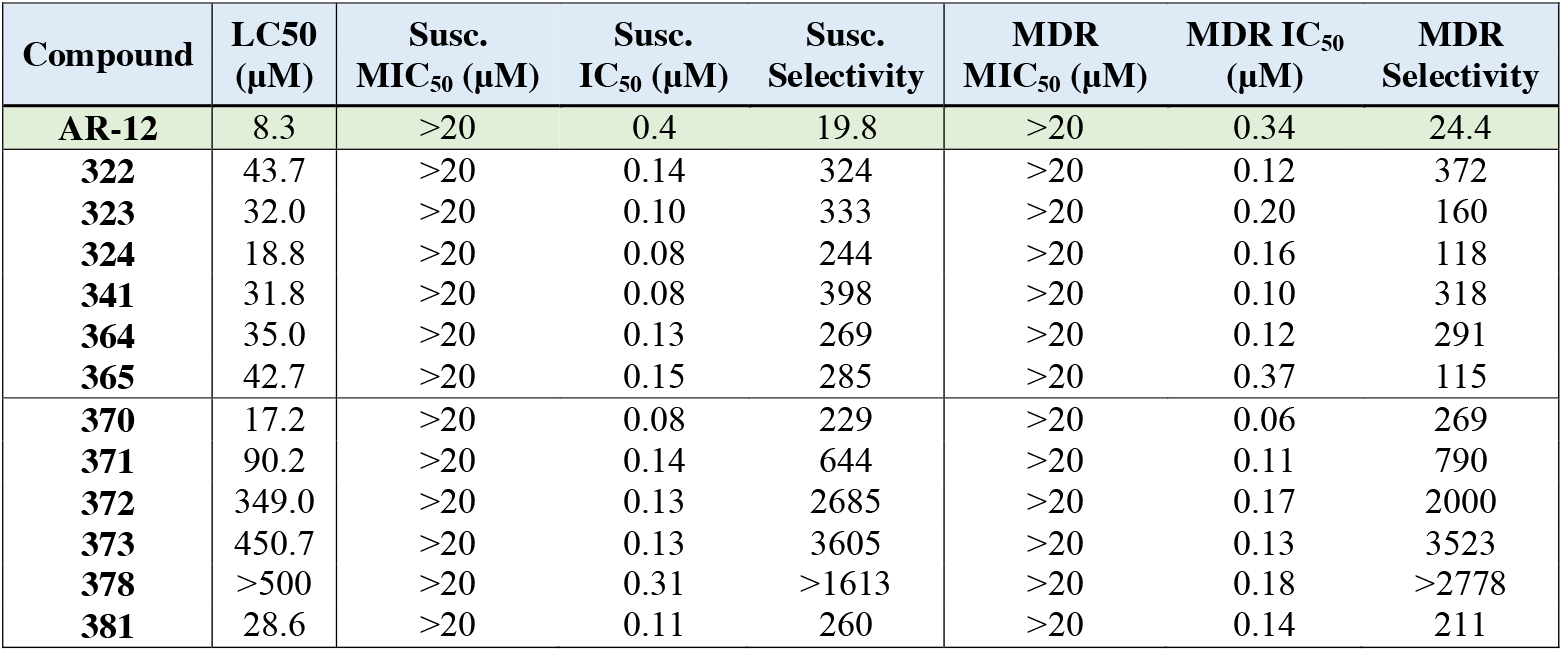
Comparing lead compounds efficacy against Susceptible and MDR *S*. Typhimurium. Concentration where RAW 264.7 macrophage cell viability is 50% (LC_50_) as determined by colorimetric assay. The minimum inhibitory concentration (MIC_50_) for each compound was determined by evaluating its inhibitory effect on planktonic *Salmonella*, using optical density measurements at 600 nm. Concentration at which intracellular *S*. Typhimurium burden is 50% in RAW 264.7 macrophages determined by gentamicin protection assay (IC_50_). Selectivity of each compound calculated by (LC_50_ / IC_50_). Parental compound AR-12 included for comparison. Susceptible (Susc.) and multi-drug resistant (MDR).

| Compound | LC50 (μM) | Susc. MIC<sub>50</sub> (μM) | Susc. IC<sub>50</sub> (μM) | Susc. Selectivity | MDR MIC<sub>50</sub> (μM) | MDR IC<sub>50</sub> (μM) | MDR Selectivity |
| --- | --- | --- | --- | --- | --- | --- | --- |
| AR-12 | 8.3 | >20 | 0.4 | 19.8 | >20 | 0.34 | 24.4 |
| 322 | 43.7 | >20 | 0.14 | 324 | >20 | 0.12 | 372 |
| 323 | 32.0 | >20 | 0.10 | 333 | >20 | 0.20 | 160 |
| 324 | 18.8 | >20 | 0.08 | 244 | >20 | 0.16 | 118 |
| 341 | 31.8 | >20 | 0.08 | 398 | >20 | 0.10 | 318 |
| 364 | 35.0 | >20 | 0.13 | 269 | >20 | 0.12 | 291 |
| 365 | 42.7 | >20 | 0.15 | 285 | >20 | 0.37 | 115 |
| 370 | 17.2 | >20 | 0.08 | 229 | >20 | 0.06 | 269 |
| 371 | 90.2 | >20 | 0.14 | 644 | >20 | 0.11 | 790 |
| 372 | 349.0 | >20 | 0.13 | 2685 | >20 | 0.17 | 2000 |
| 373 | 450.7 | >20 | 0.13 | 3605 | >20 | 0.13 | 3523 |
| 378 | >500 | >20 | 0.31 | >1613 | >20 | 0.18 | >2778 |
| 381 | 28.6 | >20 | 0.11 | 260 | >20 | 0.14 | 211 |

Evaluating these twelve compounds by chemical structure, all hits maintained a cyclohexyl group at R□, focusing diversity at R□ to modulate potency, cytotoxicity, and selectivity. Within the 4-amino piperidine series (324, 381, 322, 323), N-benzylation generally improved intracellular activity, with further gains observed when electron-withdrawing groups were present on the benzyl ring, 3-fluoro (322) and 4-cyano (323), likely enhancing lipophilicity and cell penetration. The 3-alkoxypiperidine analogs (370–373) showed a similar trend: unsubstituted derivatives exhibited strong activity, N-benzylation improved potency, and substitution with 3-fluoro (372) or 4-cyano (373) groups gave the highest selectivity indices (>2000) without loss of potent IC_50_ values. The pyrrolidine derivatives (364, 365, 378) also benefited from benzylation, with the 4-cyano benzyl analog 365 showing enhanced potency consistent with the electron-withdrawing substitution trend. Notably, glycinamide-containing analog (341) was one of the most potent at reducing intracellular bacterial burden in both susceptible and MDR strains, highlighting the potential of polar, hydrogen-bonding R□ groups, though its lower LC_50_ limited selectivity.

Compounds 372 and 373 were the most selective in the series, with IC_50_ values of ∼0.13 µM against both susceptible and MDR strains and MIC_50_ values above 20 µM, indicating a predominantly host-directed mechanism. This represents a 100-fold improvement in selectivity over AR-12 which has a selectivity near 20. While improving selectivity was a primary goal, potency against intracellular *S*. Typhimurium remained critical.

Compounds 341 and 370 were the most potent against MDR *Salmonella* and both demonstrated strong host-directed effects as evidenced by high MDR selectivity >250. However, the selectivity could be improved by reducing cytotoxicity through structural modification or formulation to improve its therapeutic index. The consistently high MIC_50_ values across the series reinforce that the primary antibacterial effect is host-mediated rather than direct bacterial killing. Future work could combine these optimized chemotypes with targeted delivery strategies to further improve in vivo efficacy and tolerability. Alternatively, formulation may improve potency and reduce toxicity. We have previously formulated AR-12 into acetalated dextran microparticles, an acid-sensitive biopolymer based on dextran that had triggered release in the intracellular compartment of the macrophage.(14, 25, 28, 30, 36) Future work can focus on in vivo evaluation of lead analogs as well as formulation for more apt delivery.

### Proteomic analysis

To evaluate potential host proteins targeted by lead compounds 370 and 341, affinity capture was performed. 370 and 341 were chemically modified (**Supplemental Figure 3A-B**) and conjugated to agarose beads using a 5-carbon spacer (termed ‘370-bead’ or ‘341-bead’). Of note, we confirmed that this chemical modification did not affect potency (**Supplemental Figure 3C-D**). We also prepared a bead conjugated to butylamine to serve as a control for nonspecific interactions (termed ‘control-bead’). Bound proteins were then evaluated by LC-MS/MS. This experiment was performed twice for 370 and once for 341. For 370 n=1,254 proteins and n=354 proteins were found to be enriched on the 370-beads compared to control beads for experiments 1 and 2, respectively. For 341 n=316 proteins were enriched on the 341-beads compared to control beads. These enriched proteins all had a Log2 fold-change >2 and p < 0.05 (**Supplemental Figure 4A)**.

Comparing the significant proteins in each experiment revealed 80 overlapping proteins in all three proteomic analyses (**Supplemental Table 3)**. Cross-referencing these proteins against the Contaminant Repository for Affinity Purification Mass Spectrometry Data (CRAPome)(37) led us to remove 13 proteins (**Supplemental Table 4**) which had average spectral counts greater than five.(37) The resulting 67 proteins were entered into Search Tool for the Retrieval of Interacting Genes/Proteins (STRING) to assess protein-protein interactions (**Supplemental Figure 4B)**. This analysis revealed a high enrichment in proteins involved in vesicle-mediated transport (**Figure 4A, Supplemental Figure 4C**). Specifically, there was strong enrichment of the Golgi transport complex and vesicle tethering complex cellular components (**Figure 4B**) which correspond to retrograde transport at the trans-Golgi-network and intra-Golgi traffic identified by Reactome Pathway Database analysis (**Supplemental Figure 4D**). We interpret this enrichment of Golgi transport and vesicle tethering complexes as likely reflecting capture of intact trafficking assemblies rather than direct interaction with all identified proteins. Especially given the highly interconnected nature of the Golgi where tethering and retrograde transport machinery exist as highly interconnected assemblies. This may be further influenced by *Salmonella* infection, where remodeling of host membrane trafficking pathways may further increase the abundance or accessibility of these complexes, enhancing their recovery.

**Figure 4:**
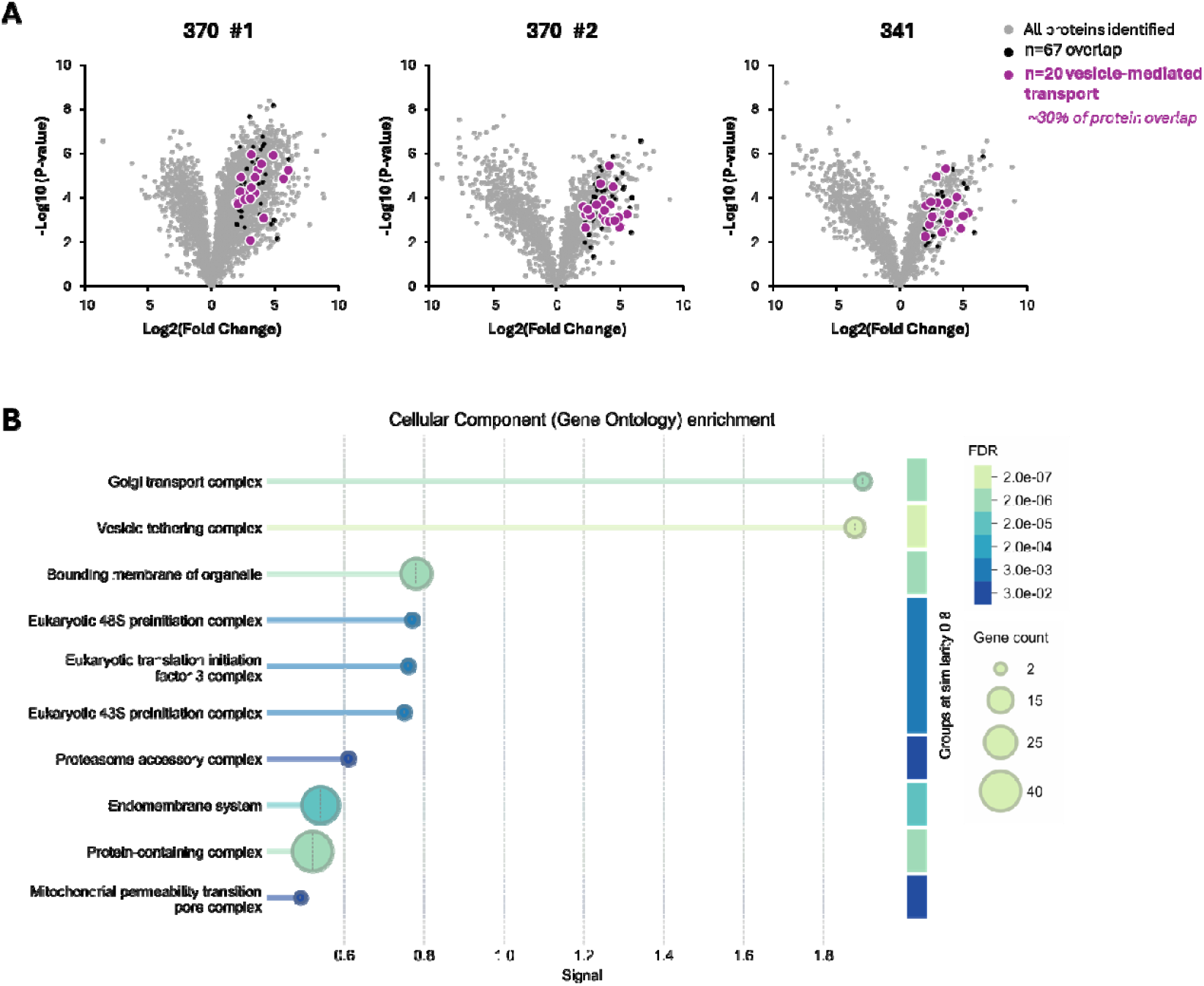
Proteomic analysis of lead compounds. **A)** Volcano plots illustrating significant proteins identified by affinity capture experiments, highlighting those proteins involved in vesicle-mediated transport. **B)** Cellular component enrichment identified by STRING database.

Intracellular *Salmonella* replication occurs in membrane bound compartments called *Salmonella*-containing vacuoles (SCV). These SCV migrate to the perinuclear region in close association with the Golgi, and importantly, an intact Golgi is essential for maintenance of SCV.(38, 39) *Salmonella* then rapidly remodels the host cell endosomal system, establishing *Salmonella*-induced filaments (SIF) to form its extensive tubular network.(40) Kehl et al. found that intracellular *Salmonella* deploys the entire late endo-/lysosomal vesicle fusion machinery to form this network, with high ranking trafficome hits including the retromer, exocyst complex, and SNAREs.(40) Through proteomic analysis, Reuter et al. found that *Salmonella* coopts conserved trafficking routes in both endothelial and macrophage cells including retromer-mediated protein sorting system, which controls the endosome to Golgi retrieval pathway, and the COPI-mediated transport, which is responsible for vesicle transport between cis-Golgi back to the endoplasmic reticulum and endosomal-lysosomal trafficking.(41) Additionally, they found that 58% of *Salmonella*-modified proteins were part of vesicles.(41) These studies make clear that intracellular *Salmonella* replication is dependent on and exploits the host cell vesicle-mediated transport system. Therefore, it is plausible that our leads compounds disrupt this fragile balance leading to a host-cell-modification that reduces intracellular *Salmonella* burden. The role of vesicle-mediated transport as a target for host-directed therapy is further supported by Singhawani et al. where knockdown of the HOmotypic fusion and Protein Sorting (HOPS) complex, a protein complex that mediates vesicle tethering and fusion in the endocytic pathway, resulted in SCV fragmentation and significantly reduced intracellular *Salmonella* replication.(42)

When we look more closely at the vesicle-mediated proteins enriched by proteomic analysis we find components of the retromer, exocyst, and conserved oligomeric Golgi (COG) complexes (**Figure 5A-B**). The retromer complex is involved in mediating endosome-to-trans-Golgi-network retrograde transport. Notably, the retromer complex has been found to associate with the SCV and is required for *S*. Typhimurium intracellular survival in macrophages.(43) The exocyst complex mediates tethering of post-Golgi vesicles to the plasma membrane prior to exocytosis. *Salmonella* has been shown to recruit the exocyst complex via SipC to facilitate bacterial entry into the host cell. (44, 45) The COG complex is a peripheral Golgi protein involved in tethering of vesicles at the Golgi apparatus prior to fusion. Additionally, the COG complex regulates SNARE recycling in the Golgi, facilitating retrograde transport.(46, 47) While the COG complex has not yet been shown to interact directly with intracellular *Salmonella*, disruption in COG complex dependent vesicles has been shown to fragment the Golgi ribbon,(48) which is essential for maintenance of the SCV.(39) These studies suggest that our lead compounds may reduce intracellular *Salmonella* burden by interfering with host trafficking complexes: (1) binding to the retromer complex could destabilize SCV integrity, (2) disrupting COG function could alter SNARE distribution and impair SCV-hijacked pathways, and (3) blocking the exocyst complex could prevent bacterial entry into host cells. Together these altered functions result in reduced intracellular bacterial burden. Further studies will be necessary to fully understand the mechanism of action of the analogs as they advance in development.

**Figure 5:**
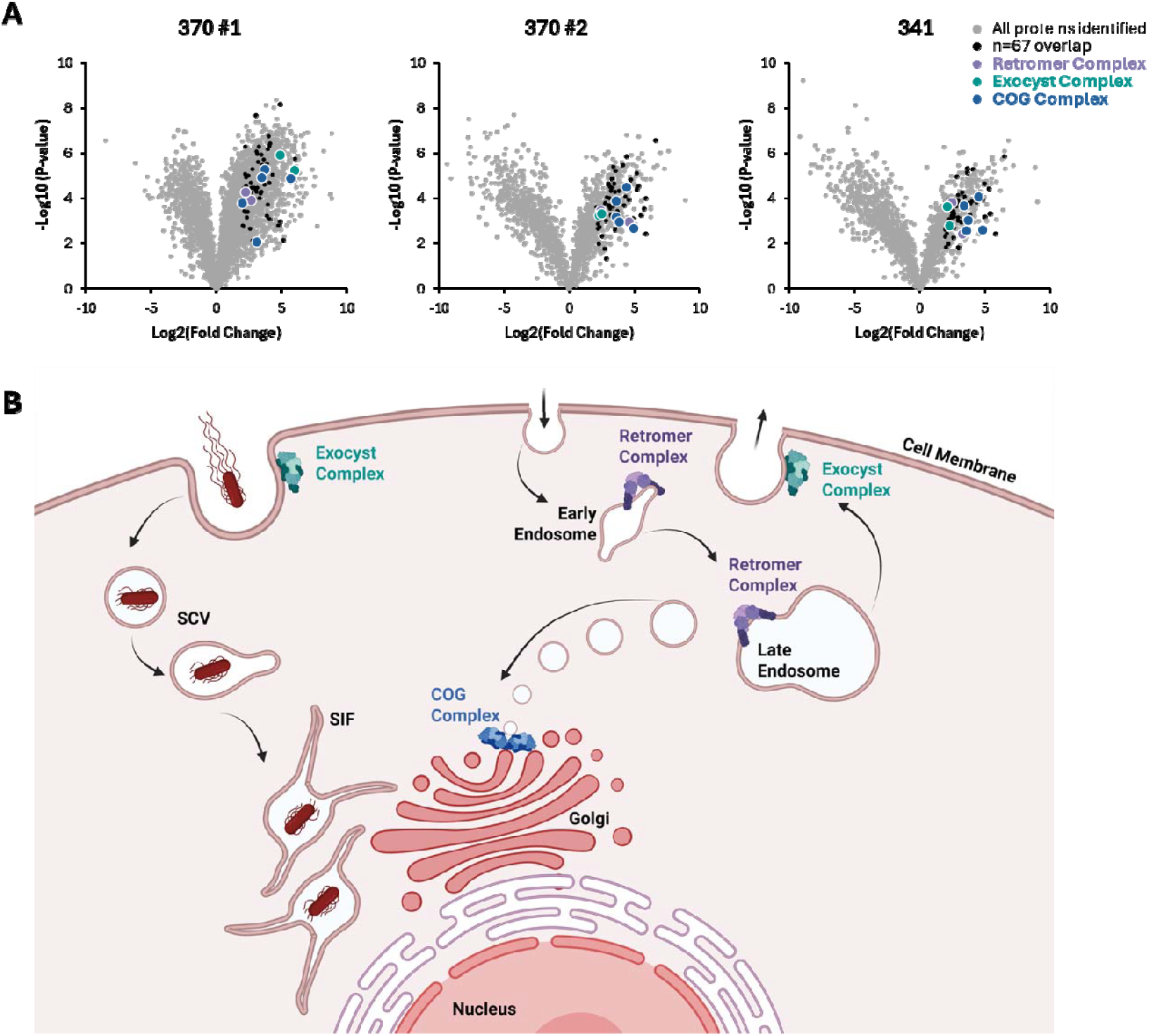
Role of vesicle tethering complexes. **A)** Volcano plots identifying significant proteins identified by affinity capture experiments, highlighting those proteins that are part of the exocyst, retromer, and conserved oligomeric Golgi (COG) complexes. **B)** Figure illustrating the overlap between vesicle-mediated transport coopted by *Salmonella* during intracellular infection and the protein complexes identified by affinity capture for compounds 370 and 341. SCV: *Salmonella*-containing vacuole, SIF: *Salmonella-*induced filament, COG: conserved oligomeric Golgi.

Salmonellosis, especially typhoid and paratyphoid fevers, significantly impacts human health around the world. Because of widespread antibiotic resistance new treatment strategies are needed. Host-targeted therapies like AR-12 show promise against various pathogens, including *Salmonella*, by leveraging host-directed mechanisms such as vesicle mediated transport. Our research has optimized AR-12 analogs for improved potency and host-directed selectivity, demonstrating significant improvements in combating MDR intracellular *S*. Typhimurium. Structure–activity analysis revealed that N-benzylation and electron-withdrawing substitutions consistently enhanced potency, selectivity, and cell penetration across multiple scaffolds, with compound 341 standing out for strong antibacterial activity despite limited selectivity. Affinity capture proteomics with lead compounds 370 and 341 identified strong enrichment in vesicle-mediated transport pathways. These results demonstrate the potential of altering host trafficking pathways as a method for host-directed therapy. Future studies should focus on in vivo evaluation and formulation for more effective delivery, aiming to enhance treatment strategies against *Salmonella* infections.

## MATERIALS AND METHODS

### AR-12 Analog Compounds

The chemical structures for all compounds can be found in **Supplemental Table 1**. Chemical synthesis was described previously. (31)

### Bacterial Strains and Mammalian Cell Lines

Susceptible (ATCC 700720) and MDR (ATCC 700408) strains of *S. enterica* serovar Typhimurium were obtained from American Type Culture Collection (ATCC Manassas, VA) and cultured in nutrient broth (BD Difco, Detroit, MA). Murine macrophage cell line RAW 264.7 (ATCC TIB-71) was maintained in Dulbecco’s modified Eagle’s medium (DMEM) (GIBCO, Invitrogen, Carlsbad, CA) supplemented with 10% fetal bovine serum (FBS) (GIBCO) and 1% penicillin-streptomycin (P/S) (GIBCO). Cells were cultured at 37°C in a 5% CO_2_ atmosphere and used up to passage 10 once received from ATCC. Susceptible and MDR activity of *S*. Typhimurium was confirmed on Mueller-Hinton agar (BD Difco, Detroit, MA) plates with antibiotic discs (Thermo Fisher Scientific, Waltham, MA) and compared to *Escherichia coli* (TOP10) and *S*. Typhimurium XEN-33 (**Table 3**).

**Table 3:** Drug sensitivity of bacterial strains. Disc inhibition zone (diameter) in mm for indicated bacteria. The experiment was performed twice where Susc. indicates susceptible and MDR is multi-drug resistant.

| Antibiotic | *E. coli*<br>(TOP10) | *S.*<br>Typhimurium<br>(XEN-33) | Susc. *S.*<br>Typhimurium<br>(ATCC700720) | MDR *S.*<br>Typhimurium<br>(ATCC700408) |
| --- | --- | --- | --- | --- |
| Streptomycin | 0 | 13 | 12 | 0 |
| Ampicillin | 17 | 18 | 19 | 0 |
| Ceftriaxo | 30 | 26 | 26 | 28 |
| Chloramphenicol | 28 | 24 | 24 | 0 |
| Ciprofloxacin | 37 | 25 | 30 | 30 |
| Nalidixic Acid | 30 | 20 | 21 | 21 |
| **Tetracycline** | 24 | 0 | 22 | 13 |
| **Kanamycin** | 20 | 0 | 17 | 17 |
| **Levofloxacin** | 34 | 27 | 32 | 32 |
| **Gentamicin** | 17 | 18 | 17 | 18 |
| **Cotrimoxazole** | 29 | 0 | 25 | 24 |

### Intracellular Gentamicin Protection Assay

RAW 264.7 cells were seeded at 3×10^4^ cells/well in a 96-well plate and maintained at 37°C for 20 hours. A single colony of susceptible or MDR *S*. Typhimurium was inoculated and cultured overnight in nutrient broth and diluted to obtain a log-phase culture stock with an adjusted optical density (OD_600_) of 0.6, equivalent to a concentration of 3×10^8^ CFU/mL (determined by a prior colony forming units (CFU) assay). The bacterial stock solution was resuspended in antibiotic-free DMEM supplemented with 2% FBS and further diluted to obtain a bacterial concentration of 3×10^6^ CFU/mL. Cell media was aspirated and replaced with 100 µL of bacteria culture in DMEM resulting in a multiplicity of infection (MOI) of 10. The cells underwent infection for two hours or 30 minutes at 37°C with susceptible or MDR bacteria, respectively. This differential infection incubation period was based on strain-specific infectivity potential. In an initial evaluation to establish MDR infection model demonstrated the MDR to be highly infective with a time-course experiment showing that >30 mins infection lysed the macrophages. Media was aspirated from the wells and replaced with DMEM (2% FBS) containing 100 µg/mL gentamicin for one hour at 37°C to eliminate extracellular bacteria. Following this step, cells were washed once with media and treated for 22 hours with analogs at varying concentrations (0.1 - 5 µM) in DMEM (2% FBS, no pen/strep) containing 10 µg/mL gentamicin to suppress extracellular bacterial growth. AR-12 treated, uninfected, and DMSO-vehicle treated cells were included as controls. Post treatment, cells were washed with PBS and lysed with 0.1% Triton-X100, a concentration sufficient to disrupt host cells while preserving bacterial integrity. The lysates containing intracellular bacteria were serially diluted and enumerated by using drop-plate assay as described previously.(27) Agar plates were incubated overnight at 37°C before CFU from each dilution were counted. Fifty percent inhibitory concentration (IC_50_) was calculated as the drug concentration that resulted in 50% reduction of intracellular bacterial burden obtained by curve fitting normalized CFU against increasing drug concentration.

### Cell Proliferation and Viability Assay

Cell proliferation and viability in the presence of the analogs was tested in the macrophage cell line RAW 264.7 using a 3-(4,5-dimethylthiazol-2-yl)-2,5-diphenyl-2H-tetrazolium bromide (MTT) assay. Cells were seeded at 3×10^4^ cells/well in 96 well plates in DMEM (10% FBS, 1% pen/strep). After an overnight incubation at 37°C, cells were treated with analogs at concentrations ranging between 1-100 µM at 37°C for 24 hr. DMSO vehicle was utilized as a control, as all the drugs were reconstituted from DMSO stocks. After 24 hr treatment, media was replaced with 100 µL of 0.6 mg/mL MTT solution and the cells were incubated at 37°C until formazan crystals formed. The MTT solution was then replaced with 100 µL of isopropanol, to dissolve the formazan crystals. Cell viability was determined by measuring absorbance at 560 nm of drug treated cells normalized to DMSO vehicle controls. The absorbance at 670 nm was used as a background correction. Cell viability was plotted against increasing drug concentration to obtain the 50% lethal concentration (LC_50_), defined as the drug concentration resulting in 50% cell viability.

### Direct Effect on Bacteria

A single colony of susceptible or MDR bacteria was inoculated into nutrient broth and incubated overnight at 37°C in an orbital shaker set at 200 rpm. The overnight culture was sub-cultured (1:100) in fresh medium to obtain log-phase bacterial growth after an additional 3 hr incubation at 37°C and further adjusted to obtain a bacterial stock solution with an absorbance value of 0.6 at 600 nm, equivalent to a concentration of 3×10^8^ CFU/mL. The bacterial stock was diluted with nutrient broth to obtain 3×10^4^ CFU/well in a 96 well plate. Next, 10 µL of each analog were added to yield a total volume of 100 µL /well with a final concentration of 20 µM. Drug-free and bacteria-free wells were used as controls. The plate was incubated at 37°C for 22 hr. Percent bacterial viability was determined using optical density measurements at 600 nm, subtracting out bacteria free absorbance (background) and normalizing to a DMSO vehicle control. Bacterial viability was plotted against increasing drug concentration to obtain the 50% minimum inhibitory concentration (MIC_50_), defined as the drug concentration resulting in 50% bacterial viability.

### Proteomic Analysis via Affinity Capture

To generate cell lysate, five million RAW 264.7 cells were plated overnight in a 10 cm dish. A single colony of susceptible *S*. Typhimurium was inoculated and cultured overnight in nutrient broth and diluted to obtain a log-phase culture stock with an adjusted optical density of 0.6, equivalent to a concentration of 3×10^8^ CFU/mL. The bacterial stock solution was resuspended in antibiotic-free DMEM supplemented with 2% FBS and further diluted to obtain a bacterial concentration of 5×10^6^ CFU/mL. Cell media was aspirated and replaced with 10 mL of bacteria culture in DMEM resulting in an MOI of 10. The cells underwent infection for 2 hr at 37°C. Media was aspirated from the wells and replaced with DMEM (2% FBS) containing 100 µg/ml gentamicin for one hour at 37°C to eliminate extracellular bacteria. Cells were washed once with PBS and then collected by scraping and pelleted with centrifugation. The cell pellet was resuspended in cell buffer (50 mM Tris-HCl, 150 mM NaCl, 2 mM EDTA) and subjected to five freeze-thaw cycles. Samples were centrifuged 13,000g x 15 minutes to remove cellular debris. Protein concentration of supernatant was determined by BCA assay and cell lysate was stored at -80 LJ C until further use.

To facilitate compound conjugation to agarose beads, a butylamine handle was added to compounds 370 and 341 (termed 424 and 490, respectively, **Supplemental Figure 3A-B**). Functionalized agarose beads were made as follows. NHS activated agarose beads were washed three times to remove acetone and resuspended in PBS to create a slurry. Compounds 424 or 490 (370 or 341, respectively, with ligation chemical handle) (0.3 mL, 25 mM in DMSO) and 1.7 mL PBS was added to 1 mL of agarose slurry for a final volume of 3 mL at a concentration of 2.5 mM (10% v/v DMSO) and incubated for 2 hrs at room temperature, then washed three times with PBS to remove unbound drug. Beads were then blocked with 1M Tris-HCl (pH 7.5) for 1 hr at room temperature. Beads were washed twice with buffer (100 mM Tris-HCl, 300 mM NaCl, 2 mM EDTA, 0.5% NP-40) and resuspended in 1 mL of buffer. Control beads were generated by coupling 2.5mM butylamine to NHS-agarose beads in the same manner as above.

400-500 µg cell lysate was incubated with 200 µL slurry at a final volume of 1 mL for 1 hr. Unbound protein was removed through four 15-minute washes. Beads were then boiled in Laemli sample buffer and resolved in a polyacrylamide gel and stained with Coomassie. Gel lanes were then excised and destained. Samples were reduced with dithiothreitol, alkylated with iodoacetamide, then digested with trypsin overnight at 37 □ C. Peptides were extracted, acidified, desalted using C18 spin columns, and analyzed by LC-MS/MS using Thermo Easy nLC 1200-QExactive HF. Data was processed using Proteome Discoverer 2.5 against Uniprot mus musculus and *Salmonella typhimurium* databases. Only proteins with > 1 peptide were reported. Data was imported into Perseus for imputation and statistical analysis. Log2(fold-change) compared to butylamine bead was plotted vs -Log10(p-value) to identify outliers. Proteins hits were identified as those with a Log2(fold-change) > 2 and p-value < 0.05.

## ACKNOWLEDGMENTS

This work was supported by grant R01AI125147.

## AUTHOR CONTRIBUTIONS

Initial Manuscript writing was Gurysh, Varma, Zahid, and Ainslie. Bacteria and cell assays were performed by Gurysh, Zahid, Johnson, Varma, Woodring and Vath. Host cell viability assays were performed in part by Pino. Figure generation and editing was performed by Hendricksen and Gurysh. Landavazo, Namjoshi, Wilson and Blough performed chemical synthesis and characterization. Bachelder, Ainslie, Gurysh, Johnson and Zahid contributed to experimental design. Supervision was performed by Bachelder, Blough and Ainslie. Funding was provided by Blough and Ainslie.

## SUPPLEMENTAL INFORMATION CAPTIONS

**Supplemental Table 1**. Chemical structures of AR-12 analogs screened for host-directed therapy against intracellular *Salmonella* infection. Parent compound AR-12 provided for reference.

**Supplemental Table 2. Compounds with direct activity against planktonic *Salmonella***. Concentration at which intracellular susceptible *S*. Typhimurium burden is 50% in RAW 264.7 macrophages determined by gentamicin protection assay (Susc. IC_50_) and concentration where RAW 264.7 macrophage cell viability is reduced by 50% (LC_50_) as determined by MTT assay. The concentration of compounds that reduce planktonic susceptible *S*. Typhimurium viability to 50% as measured by optical density (MIC_50_). Selectivity of each compound calculated by LC_50_ / IC_50_. Host-directed therapeutic (HDT) ratio calculated by MIC_50_ / IC_50_.

**Supplemental Figure 1**. Venn diagram demonstrating n=81 compound potency against intracellular susceptible *S*. Typhimurium and cytotoxicity against RAW 264.7 host cell relative to parental compound AR-12.

**Supplemental Figure 2: Evaluating lead compound efficacy against MDR *S*. Typhimurium. A)** Schematic illustrating gentamicin protection assay. Day 0 RAW 264.7 macrophages are seeded into 96-well plate and STM culture is inoculated. Day 1: RAW 264.7 cells are infected with STM at an MOI of 10 for 30 min, washed, then treated with gentamicin for 1hr to remove extracellular bacteria. AR-12 analogs added to infected cells for 22hr incubation. Day 2: RAWs are washed, lysed, diluted and then dropped onto agar plates. Day 3: colony forming units (CFUs) counted. **B)** Dose effect curves of intracellular MDR *S*. Typhimurium viability from RAW 264.7 macrophages after 22hr treatment with hit compounds. CFUs normalized to untreated controls. Parental compound AR-12 included for comparison.

**Supplemental Figure 3. A)** Chemical structure of 424 which was conjugated to functionalized agarose bead for affinity capture proteomic analysis for 370. **B)** Chemical structure of 490 which was conjugated to functionalized agarose bead for affinity capture proteomic analysis for 341. **C-D)** Dose effect curves of intracellular susceptible *S*. Typhimurium viability from RAW 264.7 macrophages after 22hr treatment with lead compounds and their ligated counterparts. CFUs normalized to untreated controls.

**Supplemental Figure 4. A)** Volcano plots illustrating significant proteins identified by affinity capture experiments. Compound 370 was evaluated twice (two separate experiments: 370 #1 and 370 #2) and 341 was evaluated once. **B)** Volcano plots illustrating overlapping proteins identified across all three affinity capture experiments. **C)** STRING (Search Tool for the Retrieval of Interacting Genes/Proteins) network of n=67 overlapping proteins identified by affinity capture across all three experiments. **D)** Reactome pathway enrichment identified by STRING database.

**Supplemental Table 3**. Overlapping proteins (n=80) identified in all three affinity capture proteomic screens. Log2 fold change (Log2FC) compared to bead control and -Log10(p-value) for each experiment listed.

**Supplemental Table 4**. Proteins (n=13) with high average spectral counts (Ave SC) identified by Contaminant Repository for Affinity Purification Mass Spectrometry Data (CRAPome).

