## Supplemental Data for "Novel compounds derived from AR-12 that demonstrate host-directed clearance of intracellular *Salmonella enterica* Serovar Typhimurium"

Kristy M. Ainslie

Professor

Division of Pharmacoengineering and Molecular Pharmaceutics

UNC Eshelman School of Pharmacy

4012 Marsico Hall, 125 Mason Farm Road

Chapel Hill, NC 27599, United States

**Supplemental Table 1.** Chemical structures of AR-12 analogs screened for host-directed therapy against intracellular *Salmonella* infection. Parent compound AR-12 provided for reference.

| **AR--12** | **202** | **203** | **229** | **230** | **232** | **247** | **285** |
| --- | --- | --- | --- | --- | --- | --- | --- |
| 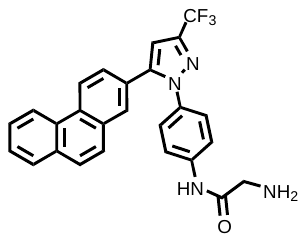 | 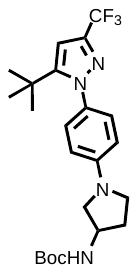 | 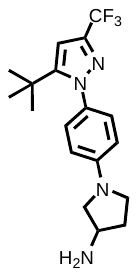 | 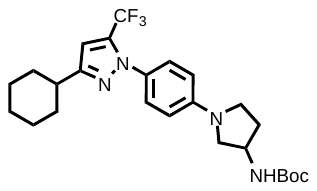 | 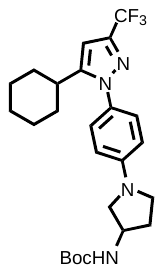 | 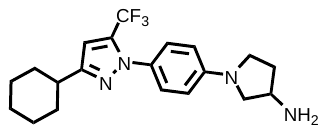 | 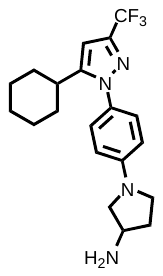 | 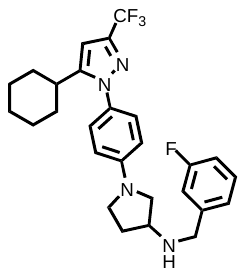 |
| **286** | **312** | **313** | **314** | **315** | **316** | **317** | **318** |
| 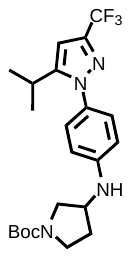 | 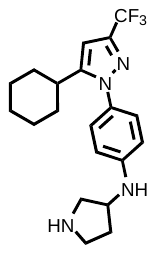 | 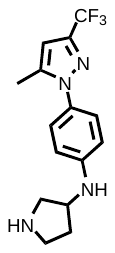 | 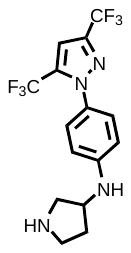 | 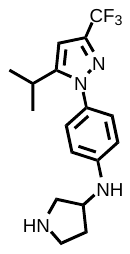 | 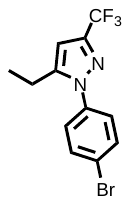 | 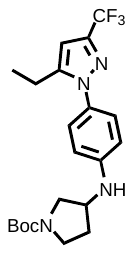 | 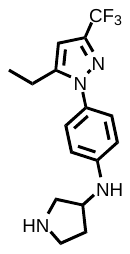 |
| **319** | **321** | **322** | **323** | **324** | **327** | **330** | **334** |
| 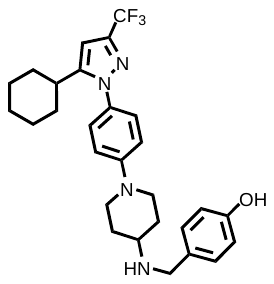 | 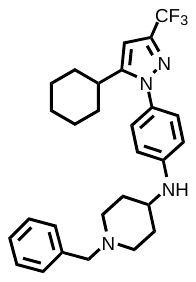 | 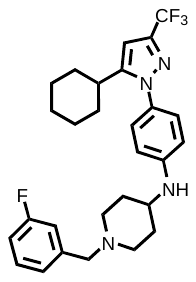 | 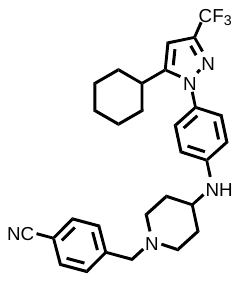 | 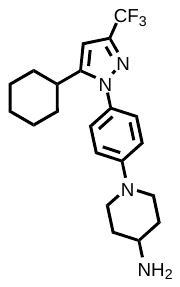 | 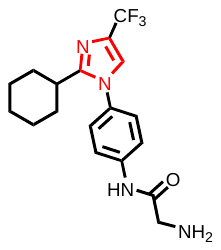 | 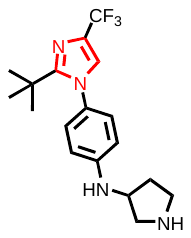 | 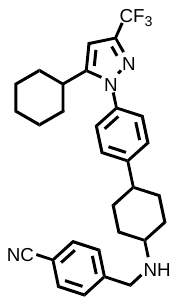 |
| **336** | **337** | **338** | **339** | **340** | **341** | **352** | **353** |
| 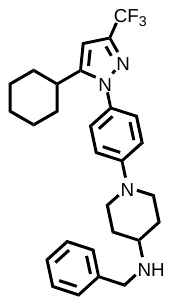 | 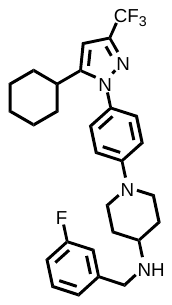 | 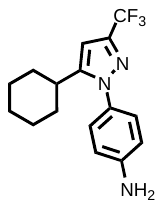 | 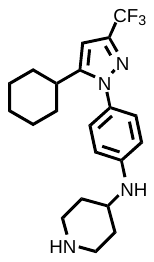 | 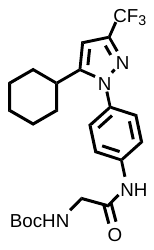 | 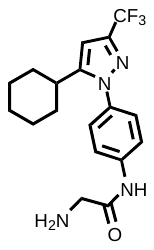 | 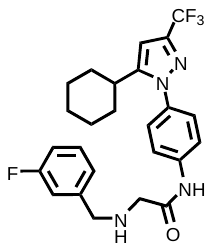 | 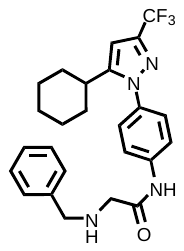 |
| **354** | **355** | **356** | **357** | **358** | **362** | **363** | **364** |
| 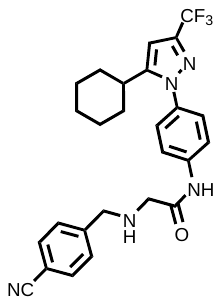 | 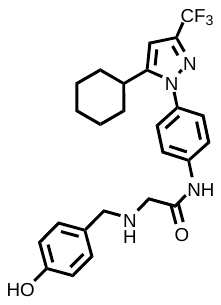 | 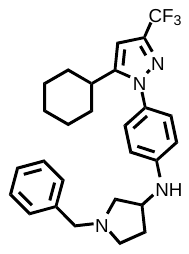 | 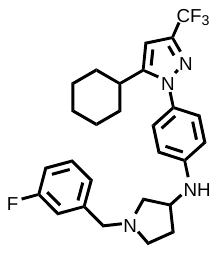 | 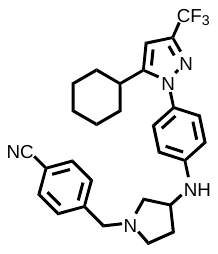 | 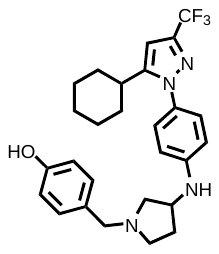 | 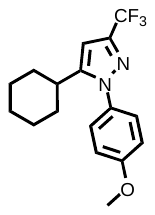 | 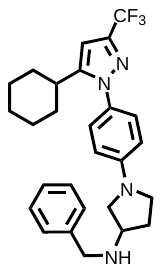 |

| **365** | **366** | **367** | **368** | **370** | **371** | **372** | **373** |
| --- | --- | --- | --- | --- | --- | --- | --- |
| 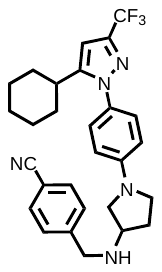 | 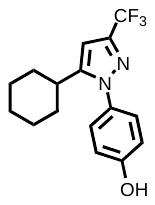 | 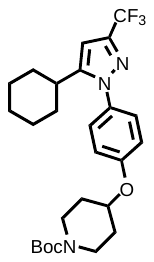 | 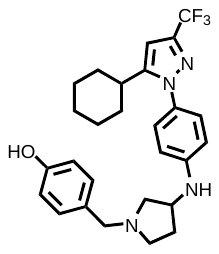 | 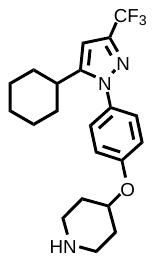 | 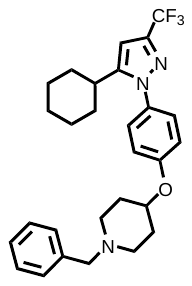 | 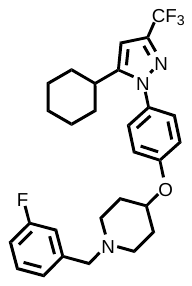 | 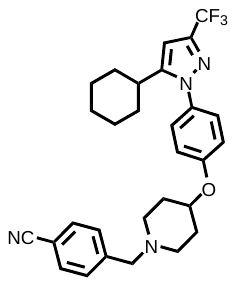 |
| **374** | **375** | **376** | **377** | **378** | **381** | **389** | **392** |
| 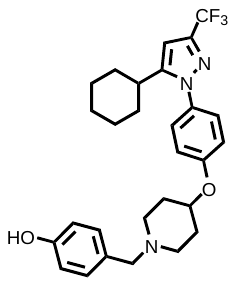 | 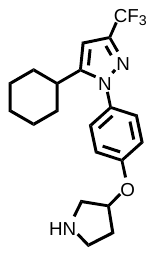 | 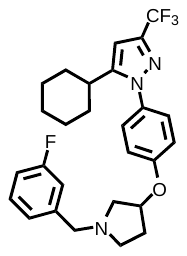 | 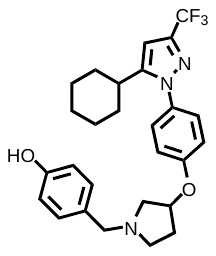 | 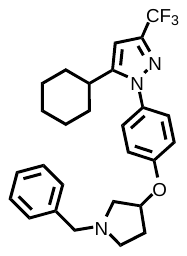 | 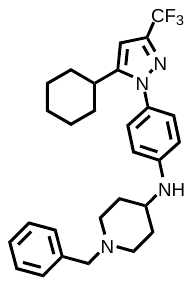 | 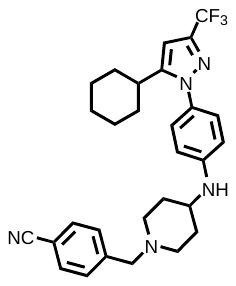 | 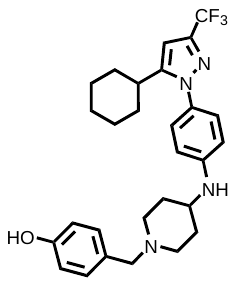 |
| **394** | **395** | **396** | **397** | **412** | **413** | **414** | **415** |
| 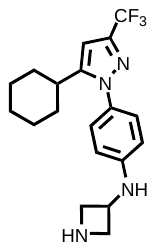 | 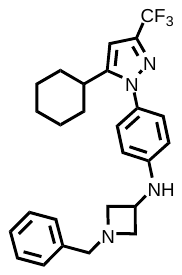 | 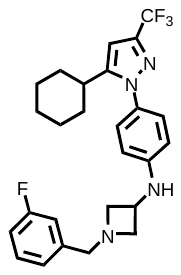 | 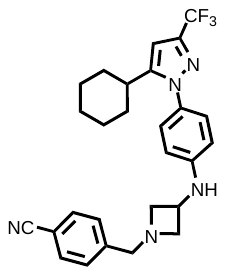 | 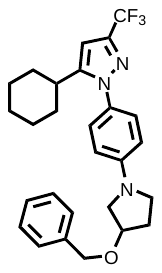 | 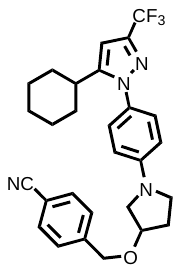 | 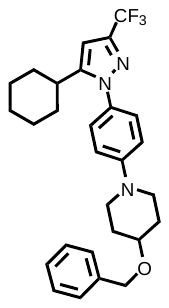 | 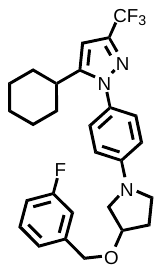 |
| **416** | **417** | **418** | **419** | **420** | **421** | **422** | **423** |
| 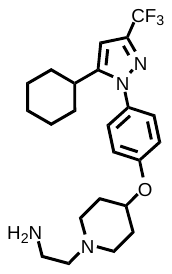 | 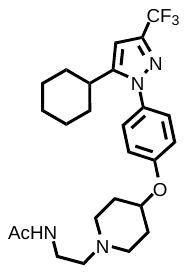 | 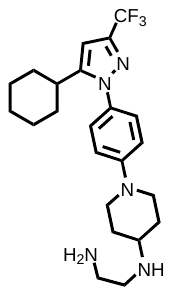 | 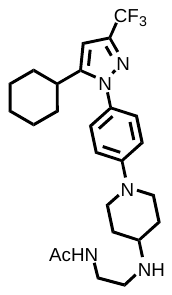 | 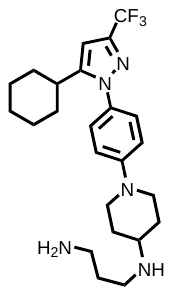 | 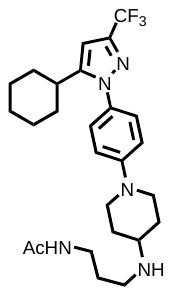 | 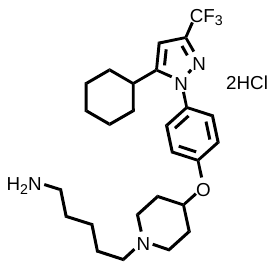 | 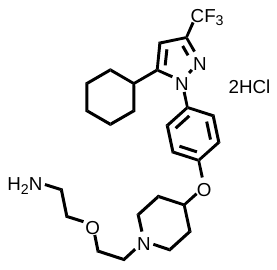 |
| **424** | **425** | **426** | **427** | **428** | **429** | **430** | **431** |
| 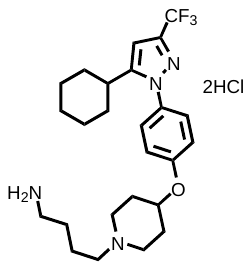 | 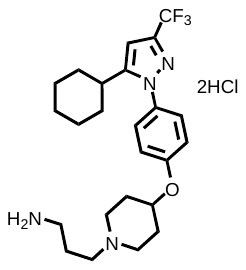 | 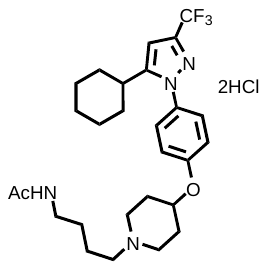 | 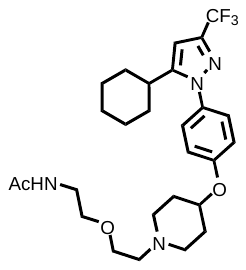 | 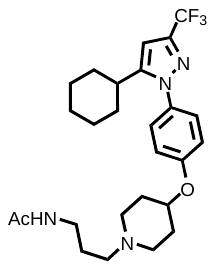 | 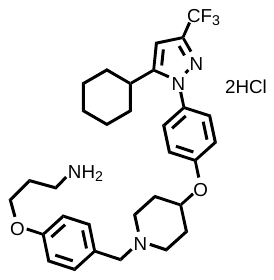 | 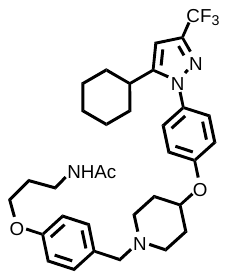 | 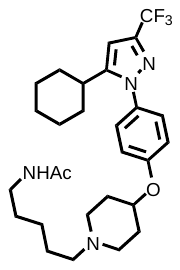 |

| **432** | **433** |
| --- | --- |
| 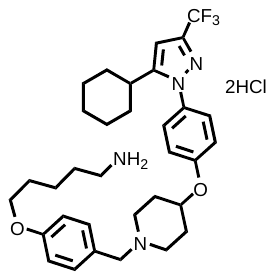 | 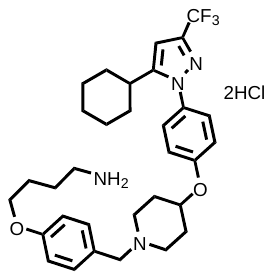 |

**Supplemental Table 2.** **Compounds with direct activity against planktonic *Salmonella*.** Concentration at which intracellular susceptible *S.* Typhimurium burden is 50% in RAW 264.7 macrophages determined by gentamicin protection assay (Susc. IC_50_) and concentration where RAW 264.7 macrophage cell viability is reduced by 50% (LC_50_) as determined by MTT assay. The concentration of compounds that reduce planktonic susceptible *S*. Typhimurium viability to 50% as measured by optical density (MIC_50_). Selectivity of each compound calculated by LC_50_ / IC_50_. Host-directed therapeutic (HDT) ratio calculated by MIC_50_ / IC_50_.

| **Compound** | **Susc.**  **IC_50_ (µM)** | **Host Cell**  **LC_50_ (µM)** | **Susc.**  **MIC_50_ (µM)** | **Selectivity (LC_50_ / IC_50_)** | **HDT Ratio**  **(MIC_50_ / IC_50_)** |
| --- | --- | --- | --- | --- | --- |
| 416 | 0.21 | 8.1 | 16.6 | 38 | 79 |
| 418 | 0.12 | 26.4 | 16.4 | 225 | 140 |
| 420 | 0.36 | 6.2 | 17.0 | 17 | 47 |
| 424 | 0.17 | 1.8 | 19.7 | 10 | 116 |
| 425 | 0.23 | 1.9 | 10.5 | 8 | 46 |
| 429 | 1.33 | 2.2 | 14.4 | 2 | 11 |
| 433 | 0.39 | 8.2 | 17.2 | 21 | 44 |

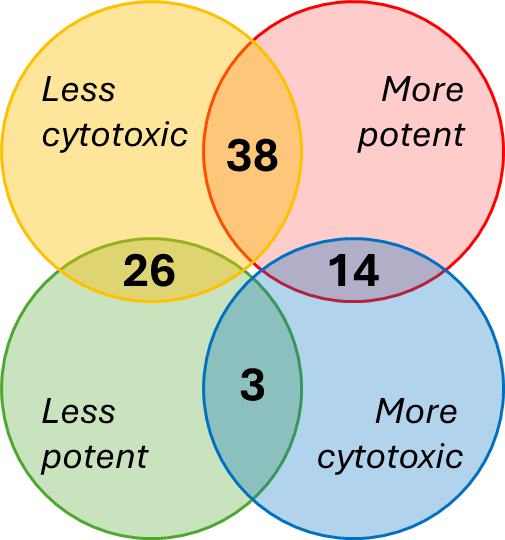

**Supplemental Figure 1.** Venn diagram demonstrating n=81 compound potency against intracellular susceptible *S.* Typhimurium and cytotoxicity against RAW 264.7 host cell relative to parental compound AR-12.

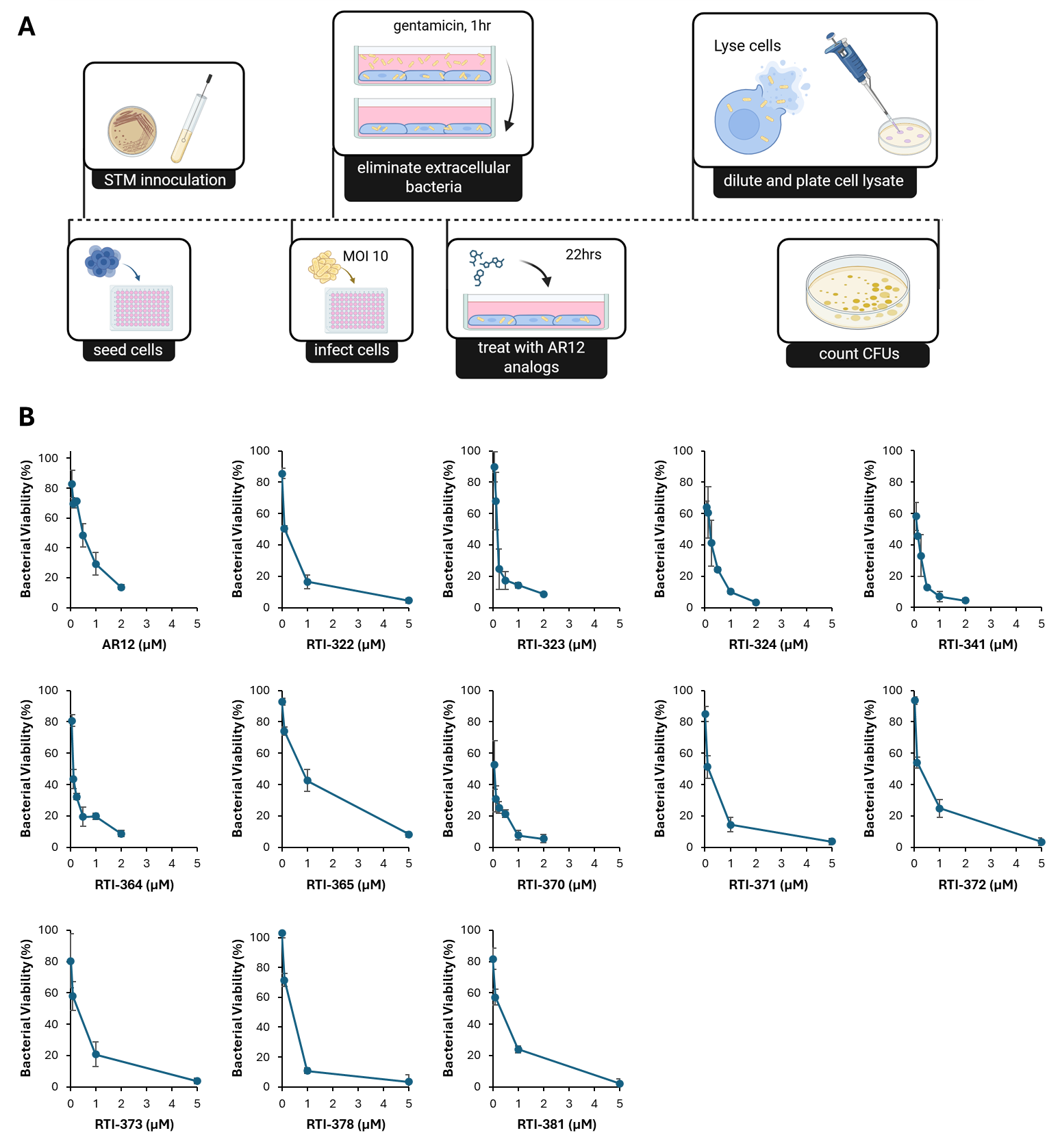

**Supplemental Figure 2: Evaluating lead compound efficacy against MDR *S*. Typhimurium**. **A)** Schematic illustrating gentamicin protection assay. Day 0 RAW 264.7 macrophages are seeded into 96-well plate and STM culture is inoculated. Day 1: RAW 264.7 cells are infected with STM at an MOI of 10 for 30 min, washed, then treated with gentamicin for 1hr to remove extracellular bacteria. AR-12 analogs added to infected cells for 22hr incubation. Day 2: RAWs are washed, lysed, diluted and then dropped onto agar plates. Day 3: colony forming units (CFUs) counted. **B)** Dose effect curves of intracellular MDR *S.* Typhimurium viability from RAW 264.7 macrophages after 22hr treatment with hit compounds. CFUs normalized to untreated controls. Parental compound AR-12 included for comparison.

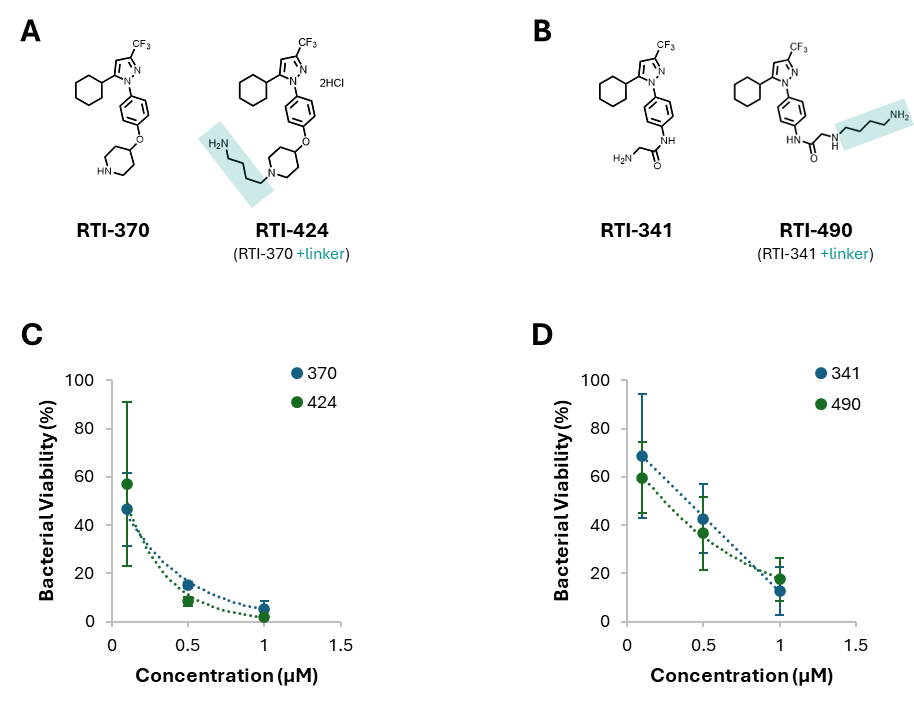

**Supplemental Figure 3.** **A)** Chemical structure of 424 which was conjugated to functionalized agarose bead for affinity capture proteomic analysis for 370. **B)** Chemical structure of 490 which was conjugated to functionalized agarose bead for affinity capture proteomic analysis for 341. **C-D)** Dose effect curves of intracellular susceptible *S.* Typhimurium viability from RAW 264.7 macrophages after 22hr treatment with lead compounds and their ligated counterparts. CFUs normalized to untreated controls.

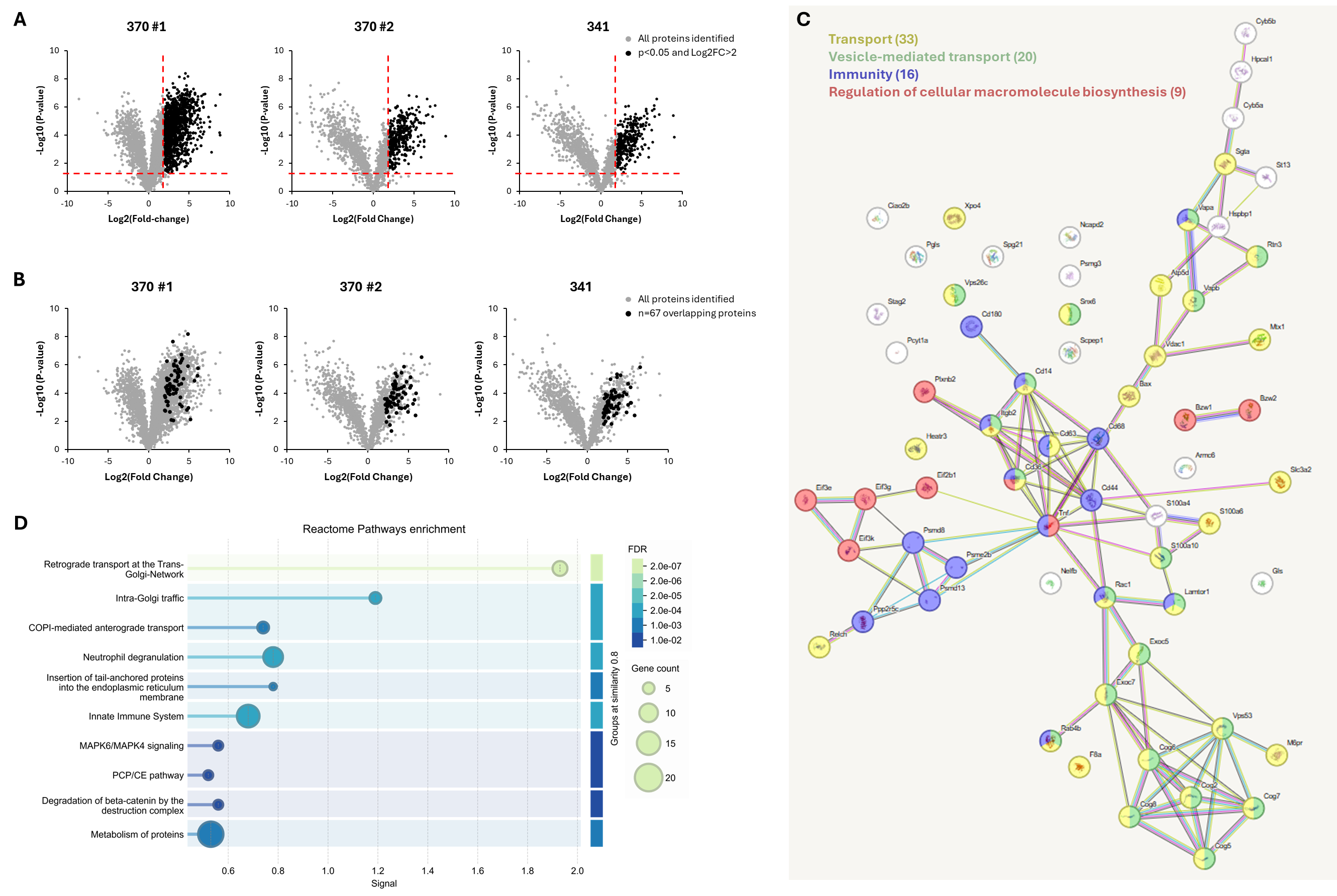

**Supplemental Figure 4.** **A)** Volcano plots illustrating significant proteins identified by affinity capture experiments. Compound 370 was evaluated twice (two separate experiments: 370 #1 and 370 #2) and 341 was evaluated once. **B)** Volcano plots illustrating overlapping proteins identified across all three affinity capture experiments. **C)** STRING (Search Tool for the Retrieval of Interacting Genes/Proteins) network of n=67 overlapping proteins identified by affinity capture across all three experiments. **D)** Reactome pathway enrichment identified by STRING database.

**Supplemental Table 3.** Overlapping proteins (n=80) identified in all three affinity capture proteomic screens. Log2 fold change (Log2FC) compared to bead control and -Log10(p-value) for each experiment listed.

|  |  | **370 #1** | | **370 #2** | | **341** | |
| --- | --- | --- | --- | --- | --- | --- | --- |
| **Accession** | **Description** | **Log2 FC** | **-Log10 (p-value)** | **Log2 FC** | **-Log10 (p-value)** | **Log2 FC** | **-Log10 (p-value)** |
| Q9CQV8 | 14-3-3 protein beta/alpha OS=Mus musculus OX=10090 GN=Ywhab PE=1 SV=3 | 6.66 | 6.29 | 2.41 | 3.15 | 2.25 | 2.12 |
| P62259 | 14-3-3 protein epsilon OS=Mus musculus OX=10090 GN=Ywhae PE=1 SV=1 | 5.91 | 6.26 | 4.00 | 3.51 | 2.69 | 3.06 |
| Q9WVJ2 | 26S proteasome non-ATPase regulatory subunit 13 OS=Mus musculus OX=10090 GN=Psmd13 PE=1 SV=1 | 4.87 | 8.18 | 2.81 | 3.33 | 2.43 | 2.91 |
| Q9CX56 | 26S proteasome non-ATPase regulatory subunit 8 OS=Mus musculus OX=10090 GN=Psmd8 PE=1 SV=2 | 3.66 | 5.71 | 3.22 | 4.30 | 2.27 | 2.85 |
| Q00558 | 40-kDa huntingtin-associated protein OS=Mus musculus OX=10090 GN=F8a1 PE=1 SV=1 | 3.10 | 4.27 | 2.72 | 1.74 | 2.98 | 1.80 |
| P10852 | 4F2 cell-surface antigen heavy chain OS=Mus musculus OX=10090 GN=Slc3a2 PE=1 SV=1 | 2.91 | 4.47 | 4.43 | 5.85 | 4.11 | 5.27 |
| Q9CQ60 | 6-phosphogluconolactonase OS=Mus musculus OX=10090 GN=Pgls PE=1 SV=1 | 3.71 | 4.60 | 3.06 | 4.62 | 2.86 | 3.64 |
| Q07813 | Apoptosis regulator BAX OS=Mus musculus OX=10090 GN=Bax PE=1 SV=1 | 3.47 | 5.92 | 3.43 | 5.39 | 3.22 | 3.71 |
| Q8BNU0 | Armadillo repeat-containing protein 6 OS=Mus musculus OX=10090 GN=Armc6 PE=1 SV=1 | 3.36 | 4.00 | 3.17 | 3.54 | 2.95 | 2.98 |
| Q91YH5 | Atlastin-3 OS=Mus musculus OX=10090 GN=Atl3 PE=1 SV=1 | 5.35 | 5.39 | 3.83 | 3.44 | 3.79 | 3.22 |
| Q9D3D9 | ATP synthase subunit delta, mitochondrial OS=Mus musculus OX=10090 GN=Atp5f1d PE=1 SV=1 | 2.39 | 3.37 | 4.11 | 4.38 | 3.65 | 5.14 |
| Q9CQC6 | Basic leucine zipper and W2 domain-containing protein 1 OS=Mus musculus OX=10090 GN=Bzw1 PE=1 SV=1 | 3.69 | 5.17 | 3.14 | 4.01 | 2.63 | 3.44 |
| Q91VK1 | Basic leucine zipper and W2 domain-containing protein 2 OS=Mus musculus OX=10090 GN=Bzw2 PE=1 SV=1 | 3.09 | 6.05 | 2.95 | 4.02 | 2.25 | 3.92 |
| P24668 | Cation-dependent mannose-6-phosphate receptor OS=Mus musculus OX=10090 GN=M6pr PE=1 SV=1 | 6.08 | 5.76 | 3.54 | 2.60 | 2.63 | 2.23 |
| Q62192 | CD180 antigen OS=Mus musculus OX=10090 GN=Cd180 PE=1 SV=2 | 3.24 | 4.87 | 5.90 | 2.41 | 5.85 | 2.41 |
| P15379 | CD44 antigen OS=Mus musculus OX=10090 GN=Cd44 PE=1 SV=3 | 2.78 | 2.10 | 4.76 | 4.81 | 4.38 | 3.82 |
| P41731 | CD63 antigen OS=Mus musculus OX=10090 GN=Cd63 PE=1 SV=2 | 2.73 | 4.46 | 5.91 | 4.00 | 4.83 | 3.92 |
| P49586 | Choline-phosphate cytidylyltransferase A OS=Mus musculus OX=10090 GN=Pcyt1a PE=1 SV=1 | 3.03 | 4.35 | 2.13 | 3.15 | 2.49 | 3.38 |
| O35638 | Cohesin subunit SA-2 OS=Mus musculus OX=10090 GN=Stag2 PE=1 SV=3 | 3.95 | 3.71 | 2.28 | 1.96 | 2.25 | 1.97 |
| Q8K2Z4 | Condensin complex subunit 1 OS=Mus musculus OX=10090 GN=Ncapd2 PE=1 SV=2 | 3.61 | 5.14 | 2.25 | 2.84 | 2.43 | 2.72 |
| Q921L5 | Conserved oligomeric Golgi complex subunit 2 OS=Mus musculus OX=10090 GN=Cog2 PE=1 SV=2 | 3.70 | 5.27 | 3.61 | 3.89 | 3.41 | 3.66 |
| Q8C0L8 | Conserved oligomeric Golgi complex subunit 5 OS=Mus musculus OX=10090 GN=Cog5 PE=1 SV=3 | 5.69 | 4.87 | 4.44 | 4.50 | 4.51 | 4.05 |
| Q8R3I3 | Conserved oligomeric Golgi complex subunit 6 OS=Mus musculus OX=10090 GN=Cog6 PE=1 SV=2 | 2.00 | 3.79 | 3.64 | 3.16 | 3.75 | 3.04 |
| Q3UM29 | Conserved oligomeric Golgi complex subunit 7 OS=Mus musculus OX=10090 GN=Cog7 PE=1 SV=1 | 3.45 | 4.92 | 4.96 | 2.67 | 4.82 | 2.61 |
| Q9JJA2 | Conserved oligomeric Golgi complex subunit 8 OS=Mus musculus OX=10090 GN=Cog8 PE=1 SV=3 | 3.08 | 2.06 | 3.87 | 2.95 | 3.59 | 2.56 |
| P56395 | Cytochrome b5 OS=Mus musculus OX=10090 GN=Cyb5a PE=1 SV=2 | 2.18 | 3.42 | 4.37 | 4.06 | 4.01 | 3.85 |
| Q9CQX2 | Cytochrome b5 type B OS=Mus musculus OX=10090 GN=Cyb5b PE=1 SV=1 | 5.15 | 2.13 | 3.60 | 5.20 | 3.01 | 4.12 |
| Q9D187 | Cytosolic iron-sulfur assembly component 2B OS=Mus musculus OX=10090 GN=Ciao2b PE=1 SV=1 | 2.47 | 4.99 | 2.88 | 1.32 | 3.54 | 3.07 |
| Q61753 | D-3-phosphoglycerate dehydrogenase OS=Mus musculus OX=10090 GN=Phgdh PE=1 SV=3 | 2.14 | 3.71 | 2.69 | 3.59 | 2.21 | 3.08 |
| Q8JZQ9 | Eukaryotic translation initiation factor 3 subunit B OS=Mus musculus OX=10090 GN=Eif3b PE=1 SV=1 | 4.52 | 6.28 | 3.00 | 6.06 | 3.04 | 6.02 |
| P60229 | Eukaryotic translation initiation factor 3 subunit E OS=Mus musculus OX=10090 GN=Eif3e PE=1 SV=1 | 4.32 | 5.03 | 3.56 | 4.27 | 3.50 | 3.74 |
| Q9Z1D1 | Eukaryotic translation initiation factor 3 subunit G OS=Mus musculus OX=10090 GN=Eif3g PE=1 SV=2 | 3.52 | 4.85 | 3.30 | 4.85 | 3.28 | 4.24 |
| Q9QZD9 | Eukaryotic translation initiation factor 3 subunit I OS=Mus musculus OX=10090 GN=Eif3i PE=1 SV=1 | 4.36 | 5.79 | 3.12 | 5.36 | 3.33 | 5.13 |
| Q9DBZ5 | Eukaryotic translation initiation factor 3 subunit K OS=Mus musculus OX=10090 GN=Eif3k PE=1 SV=1 | 4.03 | 6.72 | 3.37 | 4.47 | 2.35 | 3.75 |
| Q3TPX4 | Exocyst complex component 5 OS=Mus musculus OX=10090 GN=Exoc5 PE=1 SV=2 | 6.01 | 5.23 | 2.50 | 3.30 | 2.32 | 2.77 |
| O35250 | Exocyst complex component 7 OS=Mus musculus OX=10090 GN=Exoc7 PE=1 SV=2 | 4.87 | 5.91 | 2.28 | 3.24 | 2.10 | 3.63 |
| Q9ESJ0 | Exportin-4 OS=Mus musculus OX=10090 GN=Xpo4 PE=1 SV=2 | 2.08 | 2.74 | 5.31 | 4.51 | 5.33 | 4.42 |
| D3Z7P3 | Glutaminase kidney isoform, mitochondrial OS=Mus musculus OX=10090 GN=Gls PE=1 SV=1 | 2.24 | 6.26 | 2.83 | 3.78 | 2.65 | 2.66 |
| P01900 | H-2 class I histocompatibility antigen, D-D alpha chain OS=Mus musculus OX=10090 GN=H2-D1 PE=1 SV=1 | 2.23 | 3.40 | 2.81 | 3.73 | 2.49 | 3.20 |
| P01902 | H-2 class I histocompatibility antigen, K-D alpha chain OS=Mus musculus OX=10090 GN=H2-K1 PE=1 SV=1 | 2.01 | 2.77 | 2.32 | 3.18 | 2.14 | 2.84 |
| Q8BQM4 | HEAT repeat-containing protein 3 OS=Mus musculus OX=10090 GN=Heatr3 PE=1 SV=1 | 2.61 | 2.65 | 3.32 | 3.17 | 3.24 | 2.92 |
| Q9CQN1 | Heat shock protein 75 kDa, mitochondrial OS=Mus musculus OX=10090 GN=Trap1 PE=1 SV=1 | 2.29 | 5.19 | 2.57 | 3.82 | 2.47 | 3.49 |
| P62748 | Hippocalcin-like protein 1 OS=Mus musculus OX=10090 GN=Hpcal1 PE=1 SV=2 | 3.94 | 6.17 | 5.22 | 4.38 | 3.91 | 3.73 |
| Q99L47 | Hsc70-interacting protein OS=Mus musculus OX=10090 GN=St13 PE=1 SV=1 | 2.81 | 5.28 | 5.25 | 5.11 | 5.03 | 4.64 |
| Q99P31 | Hsp70-binding protein 1 OS=Mus musculus OX=10090 GN=Hspbp1 PE=1 SV=1 | 3.55 | 5.45 | 2.46 | 3.08 | 2.00 | 2.60 |
| Q9EPL8 | Importin-7 OS=Mus musculus OX=10090 GN=Ipo7 PE=1 SV=2 | 2.16 | 5.92 | 3.80 | 4.66 | 3.82 | 4.23 |
| P11835 | Integrin beta-2 OS=Mus musculus OX=10090 GN=Itgb2 PE=1 SV=2 | 2.65 | 4.19 | 2.27 | 2.63 | 2.02 | 2.26 |
| Q6PB66 | Leucine-rich PPR motif-containing protein, mitochondrial OS=Mus musculus OX=10090 GN=Lrpprc PE=1 SV=2 | 2.28 | 4.56 | 2.32 | 2.05 | 2.39 | 2.34 |
| P31996 | Macrosialin OS=Mus musculus OX=10090 GN=Cd68 PE=1 SV=1 | 2.34 | 4.12 | 2.32 | 3.60 | 2.24 | 2.95 |
| Q9CQC8 | Maspardin OS=Mus musculus OX=10090 GN=Spg21 PE=1 SV=1 | 2.22 | 5.06 | 2.57 | 3.42 | 2.54 | 2.77 |
| P47802 | Metaxin-1 OS=Mus musculus OX=10090 GN=Mtx1 PE=1 SV=1 | 4.06 | 4.66 | 3.84 | 4.31 | 3.46 | 4.06 |
| P10810 | Monocyte differentiation antigen CD14 OS=Mus musculus OX=10090 GN=Cd14 PE=1 SV=1 | 3.14 | 5.95 | 5.54 | 3.23 | 5.37 | 3.30 |
| Q8C4Y3 | Negative elongation factor B OS=Mus musculus OX=10090 GN=Nelfb PE=1 SV=2 | 2.37 | 3.12 | 2.38 | 3.05 | 2.12 | 2.43 |
| Q922L6 | Negative elongation factor D OS=Mus musculus OX=10090 GN=Nelfcd PE=1 SV=2 | 2.45 | 3.96 | 5.54 | 3.89 | 5.12 | 3.76 |
| Q08857 | Platelet glycoprotein 4 OS=Mus musculus OX=10090 GN=Cd36 PE=1 SV=2 | 2.42 | 3.97 | 4.91 | 3.11 | 5.02 | 3.18 |
| B2RXS4 | Plexin-B2 OS=Mus musculus OX=10090 GN=Plxnb2 PE=1 SV=1 | 3.29 | 6.29 | 3.18 | 4.48 | 2.63 | 3.28 |
| P17918 | Proliferating cell nuclear antigen OS=Mus musculus OX=10090 GN=Pcna PE=1 SV=2 | 4.70 | 6.30 | 2.32 | 3.11 | 2.20 | 2.70 |
| P97372 | Proteasome activator complex subunit 2 OS=Mus musculus OX=10090 GN=Psme2 PE=1 SV=4 | 2.28 | 4.52 | 3.14 | 3.64 | 2.97 | 3.12 |
| Q9CZH3 | Proteasome assembly chaperone 3 OS=Mus musculus OX=10090 GN=Psmg3 PE=1 SV=1 | 3.49 | 4.28 | 2.89 | 3.01 | 2.28 | 2.81 |
| P08207 | Protein S100-A10 OS=Mus musculus OX=10090 GN=S100a10 PE=1 SV=2 | 2.09 | 3.71 | 4.15 | 5.44 | 2.97 | 3.79 |
| P07091 | Protein S100-A4 OS=Mus musculus OX=10090 GN=S100a4 PE=1 SV=1 | 4.07 | 6.31 | 3.81 | 3.88 | 2.33 | 3.07 |
| P14069 | Protein S100-A6 OS=Mus musculus OX=10090 GN=S100a6 PE=1 SV=3 | 3.24 | 5.60 | 4.61 | 3.93 | 3.26 | 3.54 |
| Q148V7 | RAB11-binding protein RELCH OS=Mus musculus OX=10090 GN=Relch PE=1 SV=1 | 4.75 | 2.79 | 2.58 | 2.31 | 2.05 | 1.81 |
| Q9CQ22 | Ragulator complex protein LAMTOR1 OS=Mus musculus OX=10090 GN=Lamtor1 PE=1 SV=1 | 4.11 | 3.06 | 4.14 | 2.92 | 3.83 | 2.88 |
| P63001 | Ras-related C3 botulinum toxin substrate 1 OS=Mus musculus OX=10090 GN=Rac1 PE=1 SV=1 | 2.30 | 4.91 | 2.11 | 3.60 | 2.85 | 4.97 |
| Q91ZR1 | Ras-related protein Rab-4B OS=Mus musculus OX=10090 GN=Rab4b PE=1 SV=2 | 2.57 | 4.26 | 2.65 | 3.20 | 2.66 | 3.18 |
| Q9ES97 | Reticulon-3 OS=Mus musculus OX=10090 GN=Rtn3 PE=1 SV=2 | 3.90 | 5.52 | 3.11 | 3.69 | 2.56 | 3.13 |
| Q99P72 | Reticulon-4 OS=Mus musculus OX=10090 GN=Rtn4 PE=1 SV=2 | 4.61 | 6.53 | 2.01 | 3.85 | 2.06 | 3.35 |
| Q920A5 | Retinoid-inducible serine carboxypeptidase OS=Mus musculus OX=10090 GN=Scpep1 PE=1 SV=2 | 4.15 | 6.43 | 3.46 | 3.66 | 2.65 | 3.33 |
| Q60996 | Serine/threonine-protein phosphatase 2A 56 kDa regulatory subunit gamma isoform OS=Mus musculus OX=10090 GN=Ppp2r5c PE=1 SV=2 | 4.91 | 2.96 | 5.75 | 3.53 | 5.44 | 3.31 |
| Q76MZ3 | Serine/threonine-protein phosphatase 2A 65 kDa regulatory subunit A alpha isoform OS=Mus musculus OX=10090 GN=Ppp2r1a PE=1 SV=3 | 2.59 | 4.10 | 3.94 | 3.45 | 3.48 | 3.86 |
| Q8BJU0 | Small glutamine-rich tetratricopeptide repeat-containing protein alpha OS=Mus musculus OX=10090 GN=Sgta PE=1 SV=2 | 3.02 | 7.67 | 3.21 | 3.25 | 2.39 | 2.62 |
| Q6P8X1 | Sorting nexin-6 OS=Mus musculus OX=10090 GN=Snx6 PE=1 SV=2 | 2.23 | 4.27 | 4.58 | 2.96 | 3.30 | 2.42 |
| Q99LC8 | Translation initiation factor eIF-2B subunit alpha OS=Mus musculus OX=10090 GN=Eif2b1 PE=1 SV=1 | 2.64 | 4.67 | 3.08 | 4.59 | 2.98 | 3.89 |
| P06804 | Tumor necrosis factor OS=Mus musculus OX=10090 GN=Tnf PE=1 SV=2 | 2.10 | 4.41 | 2.38 | 2.71 | 2.38 | 2.64 |
| O35075 | Vacuolar protein sorting-associated protein 26C OS=Mus musculus OX=10090 GN=Vps26c PE=1 SV=1 | 2.65 | 3.92 | 2.43 | 3.45 | 2.47 | 3.82 |
| Q8CCB4 | Vacuolar protein sorting-associated protein 53 homolog OS=Mus musculus OX=10090 GN=Vps53 PE=1 SV=1 | 3.41 | 4.22 | 3.46 | 4.64 | 3.65 | 5.33 |
| Q9WV55 | Vesicle-associated membrane protein-associated protein A OS=Mus musculus OX=10090 GN=Vapa PE=1 SV=2 | 3.13 | 4.46 | 4.21 | 3.66 | 3.92 | 3.25 |
| Q9QY76 | Vesicle-associated membrane protein-associated protein B OS=Mus musculus OX=10090 GN=Vapb PE=1 SV=3 | 3.04 | 3.95 | 3.75 | 3.41 | 3.77 | 3.79 |
| Q60932 | Voltage-dependent anion-selective channel protein 1 OS=Mus musculus OX=10090 GN=Vdac1 PE=1 SV=3 | 3.87 | 3.22 | 6.65 | 6.57 | 6.53 | 5.84 |

**Supplemental Table 4.** Proteins (n=13) with high average spectral counts (Ave SC) identified by Contaminant Repository for Affinity Purification Mass Spectrometry Data (CRAPome).

| **Gene Symbol** | **Num of Expt. (found/total)** | **Ave SC** | **Max SC** |  | **Accession**  **Number** | **Protein**  **Description** |
| --- | --- | --- | --- | --- | --- | --- |
| LRPPRC | 249 / 716 | 11.3 | 118.0 |  | Q6PB66 | Leucine-rich PPR motif-containing protein, mitochondrial OS=Mus musculus OX=10090 GN=Lrpprc PE=1 SV=2 |
| YWHAE | 447 / 716 | 10.0 | 81.0 |  | P62259 | 14-3-3 protein epsilon OS=Mus musculus OX=10090 GN=Ywhae PE=1 SV=1 |
| TRAP1 | 478 / 716 | 7.2 | 52.0 |  | Q9CQN1 | Heat shock protein 75 kDa, mitochondrial OS=Mus musculus OX=10090 GN=Trap1 PE=1 SV=1 |
| PCNA | 268 / 716 | 6.2 | 70.0 |  | P17918 | Proliferating cell nuclear antigen OS=Mus musculus OX=10090 GN=Pcna PE=1 SV=2 |
| PHGDH | 394 / 716 | 6.2 | 42.0 |  | Q61753 | D-3-phosphoglycerate dehydrogenase OS=Mus musculus OX=10090 GN=Phgdh PE=1 SV=3 |
| EIF3I | 280 / 716 | 5.9 | 37.0 |  | Q9QZD9 | Eukaryotic translation initiation factor 3 subunit I OS=Mus musculus OX=10090 GN=Eif3i PE=1 SV=1 |
| YWHAB | 368 / 716 | 5.8 | 39.0 |  | Q9CQV8 | 14-3-3 protein beta/alpha OS=Mus musculus OX=10090 GN=Ywhab PE=1 SV=3 |
| NELFCD | 51 / 716 | 5.5 | 29.0 |  | Q922L6 | Negative elongation factor D OS=Mus musculus OX=10090 GN=Nelfcd PE=1 SV=2 |
| IPO7 | 191 / 716 | 5.3 | 19.0 |  | Q9EPL8 | Importin-7 OS=Mus musculus OX=10090 GN=Ipo7 PE=1 SV=2 |
| RTN4 | 169 / 716 | 5.1 | 36.0 |  | Q99P72 | Reticulon-4 OS=Mus musculus OX=10090 GN=Rtn4 PE=1 SV=2 |
| PPP2R1A | 316 / 716 | 5.0 | 35.0 |  | Q76MZ3 | Serine/threonine-protein phosphatase 2A 65 kDa regulatory subunit A alpha isoform OS=Mus musculus OX=10090 GN=Ppp2r1a PE=1 SV=3 |
| EIF3B | 282 / 716 | 5.0 | 41.0 |  | Q8JZQ9 | Eukaryotic translation initiation factor 3 subunit B OS=Mus musculus OX=10090 GN=Eif3b PE=1 SV=1 |
| ATL3 | 1 / 716 | 5.0 | 5.0 |  | Q91YH5 | Atlastin-3 OS=Mus musculus OX=10090 GN=Atl3 PE=1 SV=1 |
